# Tissue-wide metabolic buffering confers resilience to mitochondrial dysfunction

**DOI:** 10.64898/2026.08.27.747448

**Authors:** Stavroula Petridi, Abhilesh Dhawanjewar, Dnyanesh Dubal, Rajini Chandrasegaram, Hannah Dickmänken, Navaroshini Bala-Muraly, Bryce A. Wilson, Thomas Eve, Bryan Marzullo, Antony Hynes-Allen, Scott A. Jones, Martin S. King, Richard Butler, Marco Sciacovelli, Edmund R.S. Kunji, Daniel A. Tennant, Jelle van den Ameele

## Abstract

Mitochondria and oxidative phosphorylation (OxPhos) are essential for cellular homeostasis. However, the phenotypes caused by mitochondrial dysfunction often display remarkable tissue-specificity. What determines the susceptibility of individual cells to metabolic or mitochondrial defects in a complex *in vivo* tissue context, remains largely unknown. We find that neural stem cells (NSCs) in the developing *Drosophila* brain can maintain normal proliferation despite severe cell-autonomous OxPhos-dysfunction, provided that sufficient neighbouring cells remain metabolically intact. This tissue-wide buffering progressively fails as the proportion of NSCs with OxPhos dysfunction increases, indicating that the phenotypic threshold for mitochondrial dysfunction is an emergent property of a tissue rather than only of individual cells or cell-types. Mechanistically, we find that OxPhos-deficient NSCs activate a stress-response associated with *ATF4/crc*-transcriptional activation. NSCs upregulate lactate dehydrogenase (LDH) expression to maintain glycolysis, but their proliferation remains limited by NAD^+^ regeneration rather than by ATP production. Non-cell-autonomous rescue of NSC-proliferation depends on LDH-dependent NAD^+^-production in a brain-wide glial network connected by gap junctions and the glutamate/aspartate-transporter Eaat1. These findings demonstrate that the phenotypic threshold for mitochondrial dysfunction is determined by tissue-wide spare metabolic capacity rather than only of individual cells or cell-types. Tissue heterogeneity thus provides resilience to metabolic dysfunction, evidencing key benefits of diversity, and suggesting new therapeutic strategies to enhance endogenous metabolic buffering.

## Introduction

Tissue-specific vulnerability is a characteristic of many neurodegenerative disorders. While disease is often caused by dysfunction of ubiquitously-expressed genes or pathways, pathology only affects specific cell-types or regions within the peripheral or central nervous system (CNS). ^1^ This is exemplified by mitochondrial diseases, which are neurodegenerative conditions caused by inherited defects in mitochondrial oxidative phosphorylation (OxPhos). OxPhos is mediated by five multi-protein complexes embedded within the inner mitochondrial membrane, and serves as the principal pathway for cellular ATP production and maintaining redox homeostasis. ^2^ Although OxPhos is key to the normal function of most eukaryotic cells, mitochondrial dysfunction does not affect all cells equally. ^3^ Clinical presentation of mitochondral diseases ranges from fatal or severely-disabling childhood-conditions, to limited tissue-specific impairment in adulthood. ^4^ One of the most frequently affected tissues is the CNS, with a major impact on disability and death. ^5^ This vulnerability of the brain is likely related to its high overall energy demand, ^6^ but for reasons that remain poorly understood, even the CNS shows regional differences in susceptibility to mitochondrial dysfunction.

Much of our knowledge regarding the reliance of cells on specific metabolic pathways and energy production is based on homogeneous cell cultures or bulk tissue. This approach helps reveal broad metabolic differences between isolated cell-types or homogenised tissues ^6–10^, and how they respond to environmental changes, but largely overlooks how interactions within heterogeneous tissue-environments may shape cellular responses to disease. *In vivo*, cells reside in a specialized microenvironment that provides cell-cell interactions, signalling cues and nutrients ^11^. In cancer ^12^ or microbial communities ^13,14^, metabolic heterogeneity and cooperation between cells emerge as key mediators of therapy-resistance ^12^ and resilience to toxic stress ^15^. Tissue-wide metabolic heterogeneity may thus provide resilience to metabolic and mitochondrial dysfunction, when some cells have spare metabolic capacity ^16–18^ for non-cell-autonomous buffering of neighbouring cells.

An important source of heterogeneity within tissues is variation in cell-type composition. The CNS is mainly composed of neuronal and glial cell-types. Under normal circumstances, glial cells provide a supportive niche for neurons and neural stem cells (NSCs) ^11,19^ and are thought to convert glucose to lactate, which is secreted locally to support neuronal OxPhos ^20–22^. This division of tasks may render neurons more vulnerable to mitochondrial dysfunction than glia ^5^. Conversely, cell-type heterogeneity and metabolic cooperation may act as a buffer against mitochondrial dysfunction, with metabolic defects in individual cells compensated through spare metabolic capacity of neighbouring cells, until tissue buffering capacity becomes saturated.

Another source of heterogeneity is through genetic mosaicism, which is particularly relevant in the context of mitochondrial disease. Mitochondria contain their own genome, the mitochondrial DNA (mtDNA), which is a small circular genome that encodes 37 genes required for OxPhos and exists in many copies within each cell. mtDNA mutations are often heteroplasmic, meaning that they only affect a proportion of all mtDNA molecules in a cell, with higher heteroplasmy levels typically having a greater impact on cell function and survival. ^23–25^ High-depth sequencing and single-cell analysis have shown an extraordinary degree of mtDNA mosaicism, with substantial cell-to-cell variation in heteroplasmy levels. ^26–31^ This tissue-wide genetic heterogeneity could confer protection against mitochondrial dysfunction, when low-heteroplasmy cells provide non-cell-autonomous buffering to high-heteroplasmy neighbours.

Advanced *Drosophila* genetics, allowing combinatorial genetic manipulation of different cell types within the same tissue ^32,33^ and targeted modulation of mtDNA heteroplasmy levels ^34–36^ make *Drosophila* an excellent tractable model system to test the impact of heterogeneity on robustness when confronted with disease. Here, we find that in the developing *Drosophila* CNS, tissue-wide metabolic buffering occurs in response to genetic mitochondrial dysfunction in neural stem cells (NSCs). *Drosophila* NSCs are known to rely on OxPhos for proliferation ^37–40^. However, we observed that when OxPhos dysfunction is mosaic, through genetic inhibition of OxPhos subunits, this reliance on OxPhos inversely correlates with the number of NSCs affected by OxPhos dysfunction across the CNS. OxPhos dysfunction limits NADH oxidation and creates a redox-limited metabolic state in NSCs, that neighbouring glia can bypass through metabolic cooperation, thus creating a tissue-wide non-cell-autonomous buffering mechanism that safeguards the developing brain from mosaic mitochondrial dysfunction. Through a glia-specific RNAi-screen we found that non-cell-autonomous buffering requires glial OxPhos- and LDH-mediated NAD^+^-production and transmembrane transporters, including the aspartate/glutamate-transporter Eaat1. These glial pathways might bypass OxPhos-requirement in NSCs with mosaic OxPhos dysfunction, by providing NSCs with glial metabolites such as aspartate as an OxPhos-dependent precursor for nucleotide synthesis, to support NSC proliferation.

Our findings demonstrate that vulnerability to mitochondrial dysfunction is not solely encoded within individual cells, but also emerges from the metabolic architecture of the tissues they inhabit, offering opportunities to enhance existing buffering-capacity for future therapies.

## Results

### Mosaic mitochondrial dysfunction does not affect NSC proliferation

We previously found that NSCs in the developing *Drosophila* larval brain strictly rely on mitochondrial OxPhos to support normal proliferation and differentiation ^37–40^. Focusing on type I NSCs in the larval ventral nerve cord (VNC) ^41^, we confirmed that RNAi-mediated knockdown of nuclear-encoded subunits of Complex I of the electron transport chain (ETC) (NDUFS1-homolog, ND-75, or NDUFA10-homolog ND-42), or the ATP synthase (Complex V) subunit ATPsynO, in all NSCs slowed their proliferation rate (**Figure S1A,B**). Impaired proliferation resulted in a reduced number of NSC progeny (**Figure 1A,B**), smaller brain size (**Figure S1C**), and defects in NSC temporal patterning ^38^ and decommissioning ^38,42^ (**Figure S1D**), highlighting the broad importance of OxPhos in NSC development.

**Figure 1.**
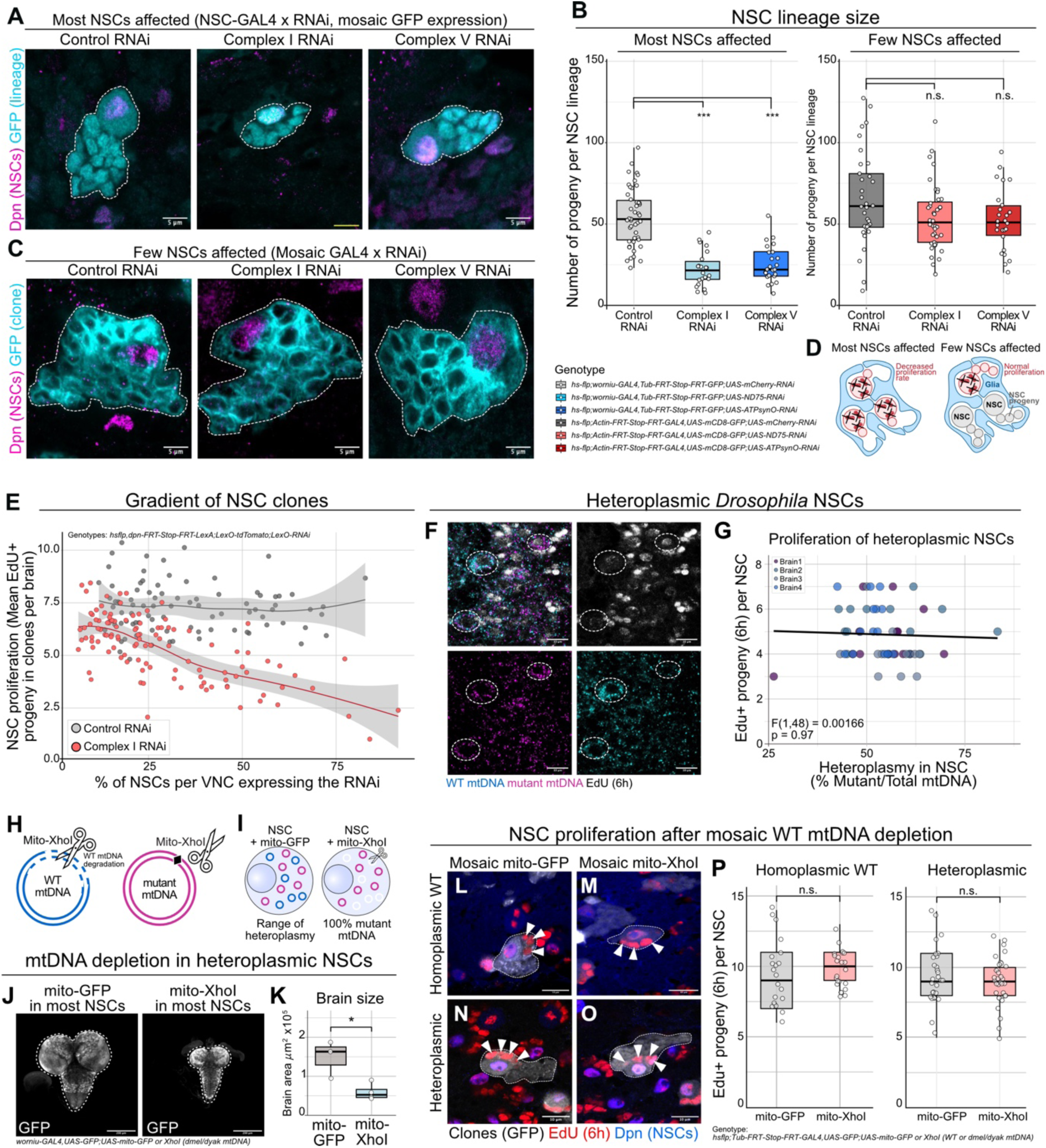
OxPhos dysfunction decreases proliferation only when most NSCs are affected. (A) NSC-lineages in VNCs from L3 larvae expressing control (UAS-mCherry RNAi), Complex I (UAS-ND75 RNAi), or Complex V (UAS-ATPSynO RNAi) RNAi in most NSCs (worniu-GAL4), with individual lineages randomly labelled by GFP, stained for Dpn (NSCs) and GFP. Dashed lines outline GFP-positive lineages from single NSCs. (B) Number of progeny per NSC lineage expressing control, Complex I, or Complex V RNAi either specifically in all NSCs or clonally; n = 46 clones, 6 brains [Most NSCs affected; Control RNAi], 22 clones, 6 brains [Most NSCs affected; Complex I RNAi], 24 clones, 5 brains [Most NSCs affected; Complex V RNAi], 33 clones, 4 brains [Few NSCs affected; Control RNAi], 40 clones, 6 brains [Few NSCs affected; Complex I RNAi], and 26 clones, 4 brains [Few NSCs affected; Complex V RNAi]; linear mixed effect models. (C) NSC-lineages in VNCs from L3 larvae expressing control, Complex I, or Complex V RNAi clonally (Actin-FRT-STOP-FRT-GAL4), with individual NSC lineages marked by GFP, stained for Dpn (NSCs) and GFP. Dashed lines outline GFP-positive lineages from single NSCs. (D) Schematic illustrating proliferation phenotypes upon reduced OxPhos function either in all NSCs or in isolated NSC lineages using mosaic approaches. (E) Number of EdU+ cells per tdTomato+ NSC lineage expressing mosaic control or Complex I RNAi in function of the percentage of NSCs expressing the RNAi defined as tdTomato+ NSCs per total Dpn+ labelled NSCs. Each dot is the average of the EdU+ progeny from 6 tdTomato+ NSCs within a single brain. The grey area around the fitted regression lines represents a 95% confidence interval. (F) mtDNA-smFISH using probes targeting only *D. melanogaster* mtDNA (mutant, magenta), or only *D.yakuba* mtDNA (WT, cyan) in L3 VNCs from heteroplasmic *Drosophila*. EdU+ progeny (grey) generated during 6h EdU feeding. Dashed outlines indicate NSCs. (G) Proliferation (number of EdU^+^ progeny) and heteroplasmy level (% mutant mtDNA determined by mtDNA-smFISH) of single NSCs in L3 VNCs; n = 14 NSCs from brain 1, 14 NSCs from brain 2, 11 NSCs from brain 3, 14 NSCs from brain 4; linear mixed effect model with brain as fixed effect. (H) Schematic of mito-XhoI activity resulting in digestion and degradation of wild-type (WT, *D. yakuba*) but not mutant (*D. melanogaster*) mtDNA. (I) Schematic of how mito-XhoI expression in NSCs increases mutant heteroplasmy levels. (J,K) Representative images (J) and quantifications (K) of brain size of heteroplasmic L3 larvae expressing control (mito-GFP) or mito-XhoI in most NSCs, labelled with GFP. Dashed lines outline whole brains. Data points indicate individual brains; n=3 brains for mitoGFP and 4 brains for mitoXhoI; unpaired t-test. (L-O) NSC lineages from L3 homoplasmic WT (L,M) or heteroplasmic (N,O) larvae clonally expressing either control (mito-GFP; L,N) or mito-XhoI (M,O). Individual NSC lineages labelled by GFP (grey), stained for Dpn (NSCs, blue) and EdU (red). Dashed lines outline GFP-positive lineages from single NSCs (Dpn+), with EdU+ progeny generated during 6hr feeding. (P) Number of EdU+ progeny of single NSCs from L3 VNCs with clonal expression (Tub-FRT-STOP-FRT-GAL4) of control (mito-GFP) or mito-XhoI in homoplasmic or heteroplasmic background. n = 53 clones, 6 brains [MitoGFP, Homoplasmic WT], n = 37 clones, 15 brains [MitoGFP, Heteroplasmic], n = 55 clones, 8 brains [MitoXhoI, Homoplasmic WT], n = 58 clones, 10 brains [MitoXhoI, Heteroplasmic]; linear mixed effect model. p > 0.05 = n.s.; p < 0.05 = *; p < 0.01 = **; p < 0.001 = ***; Scale bars: 5 μm (A,B); 10 μm (F,L-O); 200 μm (J).

We next generated ‘flip-out’ clones of NSCs, expressing either Control, Complex I or Complex V RNAis in a few NSCs only, to induce mosaic OxPhos dysfunction within otherwise normal developing larval brains. When we analysed lineage size 96 hours after clone induction, we were surprised to find that the number of progenies that were generated per NSC was not significantly different between Control NSCs and those with Complex I or V knockdown (**Figure 1B,C**). Temporal patterning was also normal, with correct repression of the early NSC temporal factor, Imp ^43^, in all clones observed (**Figure S1D-F**). This implies that, even though NSCs usually rely on OxPhos for normal development ^37–39^, proliferation and temporal patterning are not impaired when OxPhos-dysfunction is mosaic, confined to a few NSCs within otherwise metabolically healthy brains (**Figure 1D**). Using EdU-incorporation to perform lineage tracing, we quantified NSC proliferation rate in the last 6 hours before fixation, to exclude the possibility of delayed knockdown efficiency in clones versus all NSCs. While EdU-positive progeny was significantly reduced compared to control when all NSCs expressed Complex I or Complex V RNAi (**Figure S1G,I**), mosaic RNAi expression resulted in comparable numbers of EdU-positive progeny in clones expressing Control, Complex I or Complex V RNAi (**Figure S1H,J**). We validated efficacy of clonal RNAi at knocking down OxPhos subunits (**Figure S1K,L**). Moreover, similar observations were made with different genetic approaches for achieving mosaic knockdown, using different NSC-specific GAL4 ^44,45^ or LexA ^32^ driver lines (**Figure S2A,B**).

To test if the proliferation rate of affected NSCs correlated with the number of NSCs affected by OxPhos dysfunction across a brain, we gradually increased the proportion of NSCs affected by OxPhos inhibition by increasing heat-shock time at larval hatching (**Figure S2C,D**). For NSCs expressing Complex I RNAi, EdU-incorporation was inversely correlated with the number of NSC clones across the CNS, but not for NSCs expressing Control RNAi (**Figure 1E; S2E,F**). Together, this indicates the existence of a tissue-wide buffering mechanism against NSC-specific OxPhos dysfunction, which gradually becomes overwhelmed and fails when too many NSCs within the tissue are affected.

### MtDNA heteroplasmy leads to mosaicism that does not affect proliferation

To study tissue-wide buffering in a physiological and disease-relevant context of somatic metabolic mosaicism, we next turned to a *Drosophila* model of mtDNA heteroplasmy. In these flies, a temperature-sensitive lethal *D. melanogaster* mtDNA mutation (*mt:CoI^T300I^*) impairs Cytochrome Oxidase (Complex IV) activity at higher temperatures (29°C), but lethality is prevented by the co-existence of wild-type mtDNA from *D. yakuba* species within the same cells ^34,46–48^ (**Figure S3A**). We previously reported strong cell-to-cell variability in heteroplasmy levels by pyrosequencing of individual sorted NSCs ^48^, ssimilar to observations in postnatal mouse brain ^27^. However, NSC mitotic index (**Figure S3B**) and brain size (**Figure S3C**) were not significantly different between heteroplasmic flies and flies only carrying wild-type *D. melanogaster* mtDNA (homoplasmy). To determine whether the heteroplasmy levels of individual NSCs correlates with their proliferation rate, we took advantage of a recent single-molecule fluorescent in situ hybridisation (smFISH) approach to distinguish *D. melanogaster* and *D. yakuba* mtDNA *in vivo* (mtDNA-smFISH) (**Figure S3A**) ^48^. We performed mtDNA-smFISH in heteroplasmic larval CNS and found that the NSC proliferation measured by EdU incorporation did not correlate with *in situ* NSC heteroplasmy levels (**Figure 1F,G**).

Cellular dysfunction in response to mtDNA heteroplasmy may only occur above a certain threshold of mutation burden ^49^. We therefore sought to further increase NSC-specific heteroplasmy levels, by selectively depleting wild-type mtDNA, either in all NSCs across the brain, or in a mosaic manner. Wild-type *D. melanogaster* or *D. yakuba* mtDNA molecules contain an XhoI restriction enzyme recognition site, which is absent from mutant *mt:CoI^T300I^* mtDNA (**Figure 1H**) ^36,47^. NSC-specific expression of mitochondrial-targeted XhoI (mito-XhoI) in all NSCs across the larval brain results in pupal lethality ^34^, by selectively depleting wild-type mtDNA, as confirmed by mtDNA-smFISH (**Figure 1I; S3D,E**). Mito-XhoI expressing larvae have a severely reduced brain size compared to control brains expressing mitochondrial-targeted GFP (mito-GFP), both in heteroplasmic (**Figure 1J,K**) and in wild-type homoplasmic (**Figure S3F,G**) strains, indicating a requirement for mtDNA and Complex IV activity to sustain NSC proliferation. In contrast, clonal expression of mito-XhoI, to deplete wild-type mtDNA in a few NSCs only, did not decrease NSC proliferation rate compared to mito-GFP, both in homoplasmic wild-type and in heteroplasmic brains (**Figure 1L-P**). Taken together, these results indicate that the tissue-wide heterogeneity caused by cell-to-cell variability in mtDNA heteroplasmy levels allows NSCs with high mtDNA mutation burden to sustain a normal proliferation rate, provided they reside in an otherwise wild-type or heteroplasmic context. This is even the case when NSCs are fully depleted of wild-type mtDNA, or carry mutant mtDNA heteroplasmy levels above thresholds otherwise leading to dysfunction.

### NSCs compensate for OxPhos dysfunction through LDH-mediated increase of glycolysis

To understand how OxPhos dysfunction affects NSCs and their surrounding tissue, we first performed single-cell RNA sequencing (scRNAseq) on second instar larval VNCs expressing either Control or Complex I (*ND-75*) RNAi in all NSCs (**Figure 2A**). scRNAseq was conducted in two replicates, yielding 87,674 cells post-QC (44,051 Control RNAi; 43,623 ND-75 RNAi). Clustering after integration and batch-correction (**Figure S4A-C**) resulted in 19 clusters at resolution 0.4, which were visually represented using UMAP (**Figure 2B; S4D-F**). Clusters were manually annotated based on previously identified cell-type specific marker genes (**Figure 2C; S4G; Table S1**) ^50,51^ in combination with NSC-specific mCD8-GFP and glia-specific RFP transgene expression (**Figure S4H**). We focused our downstream analysis on three main cell type categories, physically and phylogenetically most closely related to NSCs: NSCs (Cluster 13); ganglion mother cells (GMCs), which are the immediate progeny of NSCs (Clusters 4, 14 and 15); and the various glial cell types surrounding the NSCs, including cortex glia (Cluster 16), surface glia (Cluster 10) and astrocytes (Cluster 7) (**Figure 2C; S4F,G)**. Differential gene expression analysis using edgeR ^52^ confirmed significant downregulation of the Complex I target gene *ND-75* in NSCs only (**Figure 2D; S5A,B; Table S2**). Module-based gene expression analysis using hdGCWNA ^53^ focusing on NSCs, identified 11 gene co-expression modules (**Figure 2E; S5C,D; Table S3**). Gene ontology analysis showed that the most strongly upregulated modules contained genes involved in stress response (M7), gene transcription (M5, M6), mitochondrial biogenesis and NAD/carbohydrate metabolism (M9), while downregulated modules were related to protein degradation (M2), chromatin regulation (M2, M8) and mitochondrial OxPhos (M1) (**Figure 2E; S5D; Table S4**). Network analysis of these gene modules showed clear function-based separation of up- and downregulated genes, with genes like *Ldh*, *Hex-A*, *Xrp1*, *Ets65a* and *Tspo* from module M9 (mitochondrial biogenesis and NAD/carbohydrate metabolism module) bridging the upregulated stress response and chromatin-regulatory gene clusters with the downregulated mitochondrial ATP-synthesis/proteostasis gene clusters (**Figure 2F; Tables S5,S6**). Intersecting differentially expressed genes detected by both edgeR and hdWGCNA in response to Complex I knockdown revealed 10 significantly differentially expressed genes (**Figure 2G; Table S7**), with lactate dehydrogenase (*Ldh*) being most strongly upregulated. LDH is a key glycolytic enzyme involved in pyruvate reduction to lactate and back. Pyruvate reduction allows regeneration of NAD^+^ from NADH to sustain glycolysis-mediated ATP production upon oxygen deprivation or OxPhos dysfunction (**Figure 2H**). Complex I knockdown-mediated *Ldh* upregulation was restricted to NSCs, as the expression levels of *Ldh* in other cell types, including GMCs and glial cell types, remained unchanged (**Figure 2I**). *Ldh* upregulation upon OxPhos dysfunction (either Complex I or Complex V knockdown) was further confirmed in NSCs *in vivo* by assessing transcription using a transgenic reporter ^54,55^ (**Figure S5E**) and LDH protein expression by immunostaining (**Figure 2J**).

**Figure 2.**
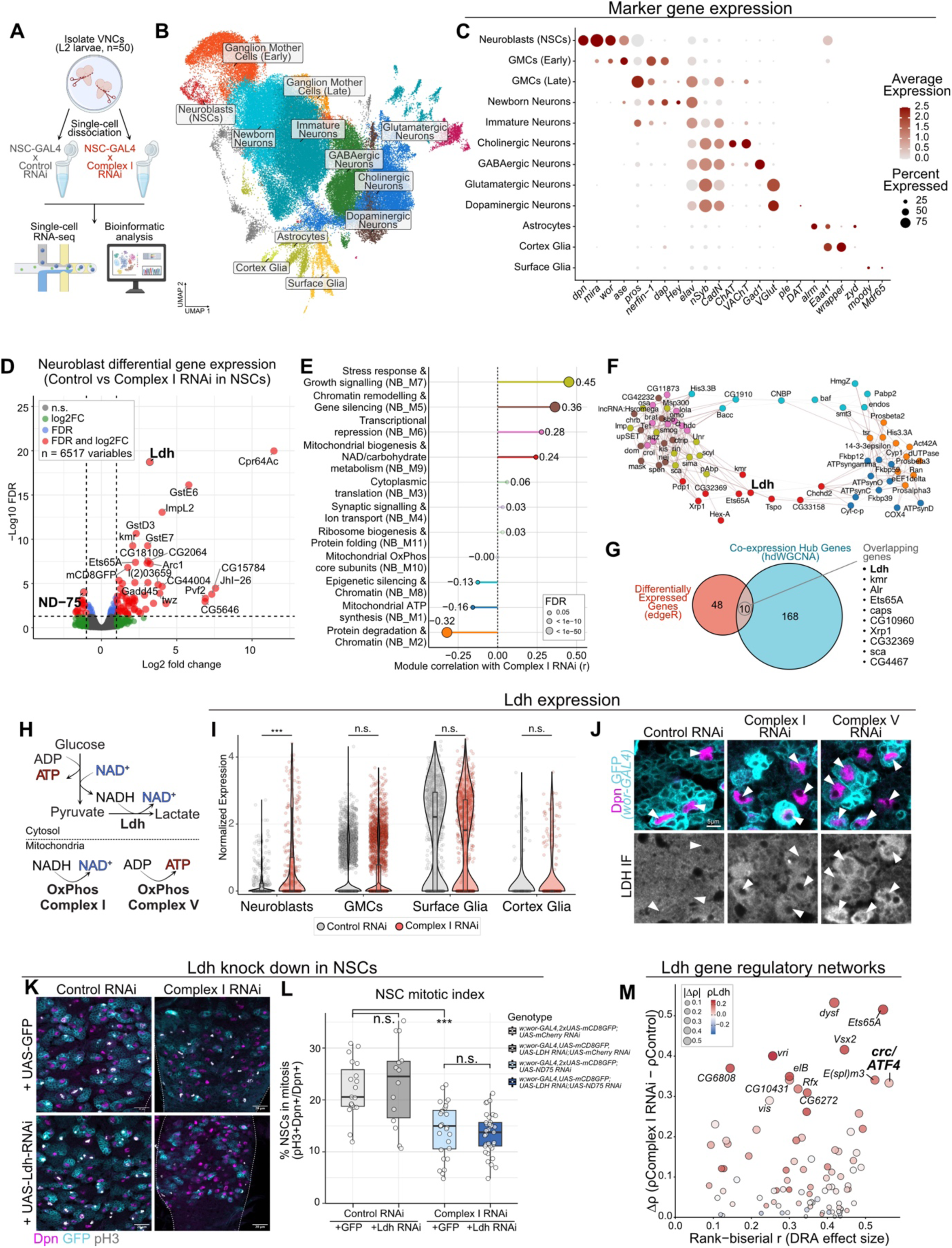
NSCs compensate for OxPhos dysfunction through LDH-mediated increase of glycolysis. (A) Experimental approach for the scRNA seq experiment from L2 VNCs. (B) UMAP of single-cell transcriptomes from L2 VNCs, colours according to cell-type annotation. (C) Expression of cell-type marker genes used to assign cell-type annotations. Dot size represents percentage of cells expressing each gene; colour intensity indicates average expression level. (D) Pseudobulk differential gene expression (edgeR) between control and mitochondrial dysfunction (Complex I RNAi) in NSCs. Significant differentially expressed genes defined as FDR < 0.05 and absolute log2 fold change ≥ 1. Colours indicate genes passing FDR threshold, log2 fold-change threshold, both thresholds, or neither; selected differentially expressed genes are labelled. (E) Association of NSC gene co-expression modules (hdWGCNA) with Complex I RNAi. The x-axis shows the correlation coefficient (r) between module eigengenes and condition; dot size represents FDR. Modules are labelled according to representative enriched biological processes identified by Gene Ontology. (F) Network visualization of hub genes from NSC gene co-expression modules (hdWGCNA) significantly associated with Complex I RNAi. Nodes represent genes, coloured by module membership; edges represent co-expression network connections. (G) Venn diagram showing overlap between pseudobulk differentially expressed genes identified by edgeR and co-expression hub genes identified by hdWGCNA in NSCs. (H) Schematic representation of cellular energy and redox metabolism linking glycolysis, lactate production, and oxidative phosphorylation. (I) *Ldh* expression levels in control versus Complex I RNAi conditions. (J) Dpn (NSCs) and LDH immunostaining in L3 VNCs expressing control (*mCherry*-RNAi), Complex I (ND75 RNAi) or Complex V (*ATPSynO* RNAi) RNAi in most NSCs (worniu-GAL4) with individual NSC lineages marked by GFP. White arrows indicate NSCs. (K) Phospho-Histone H3 (pH3), Dpn (NSCs) and GFP (NSC lineages) staining in L3 VNCs expressing control or Complex I RNAi in most NSCs (worniu-GAL4), together with either control (UAS-GFP) or UAS-Ldh-RNAi. Individual NSC lineages marked by GFP. Dashed lines outline the VNC. (L) Mitotic index of Dpn+ NSCs from L3 VNCs expressing the indicated RNAi constructs in most NSCs (worniu-GAL4). Each datapoint represents an individual brain, n = 19 [Control RNAi; GFP], 14 [Control RNAi, LDH RNAi], 28 [Complex I RNAi; GFP], 35 [Complex I RNAi; LDH RNAi]; linear mixed effect models. (M) pySCENIC analysis of transcription factor regulon activity and its association with Ldh expression in NSCs. x-axis shows the rank-biserial effect size for differential regulon activity between Complex I RNAi and Control; y-axis shows change in Spearman correlation between regulon activity and Ldh expression. Regulons with increased activity and increased Ldh coupling in Complex I RNAi are shown. Dot size represents absolute change in Spearman correlation; colour represents the overall correlation between regulon activity and Ldh expression. p > 0.05 = n.s.; p < 0.05 = *; p < 0.01 = **; p < 0.001 = ***. Scale bars: 5 μm (J); 20 μm (K).

In wild-type *Drosophila* NSCs, *Ldh* is absent or expressed at very low levels (**Figure 2I**). *Ldh* expression is not required in NSCs for their proliferation under control conditions (**Figure 2K,L**) but its overexpression caused a significant decrease in NSC proliferation (**Figure S5G,H**), further confirming previous observations that highly proliferative NSCs *in vivo* normally do not rely on aerobic glycolysis ^38^. However, upregulation of *Ldh* gene expression becomes essential in NSCs with OxPhos dysfunction, as combined RNAi-mediated knockdown of *Ldh* and Complex I, does not improve NSC mitotic index compared to Complex I knockdown alone (**Figure 2K,L**), and possibly further reduces lineage size (**Figure 2K**). The efficiency of *Ldh* RNAi-mediated knockdown was validated using a cortex glia-specific driver (**Figure S5F**).

To identify the gene regulatory networks underlying differential *Ldh* expression, we used single-cell regulatory network inference and clustering (SCENIC) analysis. ^56,57^ SCENIC identified key transcription factors involved in *Ldh* upregulation and the response to OxPhos dysfunction (**Figure S6A; Table S8**). The strongest differential gene regulatory network activity was associated with *crc*, the *Drosophila* homolog of the key mammalian mitochondrial stress-response transcription factor *ATF4* ^26,58^ (**Figure 2M; S6A-C**), which was previously found to act upstream of *Drosophila Ldh* expression in response to oxidative stress ^59,60^. In addition, several other NSC-specific transcription factors, including *Ets65a* and *E(spl)m3* ^50^, were significantly associated with differential *Ldh* expression (**Figure S6A-C**). Our scRNAseq data thus indicate that NSCs raise a strong, cell-autonomous stress response to OxPhos dysfunction, mediated by both generic and NSC-specific transcription factors, which primarily results in compensatory activation of *Ldh* expression and glycolysis.

### NSCs proliferation relies on OxPhos to maintain the redox balance

We next asked ourselves which metabolic consequences of OxPhos dysfunction result in a reduced NSC proliferation rate, reasoning that this would provide insight into the putative mechanisms underlying tissue-wide compensation in brains affected by mosaic OxPhos dysfunction. A key role for OxPhos is in ATP production. ^61^ However, AMPK phosphorylation, a sign of low cellular ATP levels ^62,63^, was not increased in VNCs where all NSCs expressed Complex I or V RNAis (**Figure S7A-C**), indicating that the effects of OxPhos dysfunction extend beyond ATP production. This confirms our previous findings using a genetically-encoded ATP-sensor ^38,64^, that glycolysis can sustain ATP production in NSCs with OxPhos dysfunction, and similar observations in the *Drosophila* eye disc ^65^. OxPhos dysfunction may also cause excess production of reactive oxygen species (ROS) ^66^, which may lead to a wide range of cellular phenotypes ^67^. Using dihydroethidium (DHE) staining as a measure for ROS levels, we saw increased ROS in NSCs with Complex I knockdown, and to a lesser extent with Complex V knockdown (**Figure S7D,E**). However, NSC-specific expression of a mitochondrial superoxide dismutase (SOD2) to reduce ROS levels ^68,69^ could not rescue proliferation of NSCs affected with Complex I or V knockdown (**Figure S7F**).

Findings in mammalian cell culture ^70–72^ and cancer ^73^ showed that a key role for OxPhos in proliferating cells *in vitro* is not ATP production, but instead for maintaining the cellular redox balance through regeneration of NAD^+^ from NADH. NAD-metabolism was indeed among the main Gene Ontology terms within the *Ldh* gene hdGCWNA co-expression module (M9; **Figure 2E**). As a measure for the NSC NADH/NAD^+^-ratio, we first assessed phosphorylation of pyruvate dehydrogenase (p-PDH), which is mediated by the NADH-responsive enzyme pyruvate dehydrogenase kinase (PDK). ^72,74,75^ NSC p-PDH levels were increased when all NSCs expressed a Complex I or Complex V RNAi (**Figure 3A**), but not upon mosaic OxPhos dysfunction (**Figure 3B**). We therefore tested whether reducing the NADH/NAD^+^ redox balance through genetic means could rescue proliferation of *in vivo* NSCs with OxPhos dysfunction. Complex V inhibition directly prevents ATP production but also increases mitochondrial membrane potential (MMP) across the inner mitochondrial membrane (IMM) with subsequent retrograde inhibition of the ETC and the NADH dehydrogenase activity of Complex I ^71,76,77^ (**Figure 3C**). Therefore, in order to separate ATP production from NAD^+^ regeneration, we genetically uncoupled the MMP in NSCs via ectopic expression of mammalian uncoupling proteins UCP1 or UCP2, together with RNAi-mediated inhibition of Complex I or V (**Figure 3C**). UCP1/2 expression increased brain size in all conditions (**Figure S7G,H**), possibly related to the key role of the NADH/NAD^+^ ratio in regulating developmental speed ^65,76,78^. However, UCP1/2-expression only rescued NSC mitotic index upon Complex V inhibition, but not Complex I inhibition (**Figure 3D,E; S7I,J**). This is compatible with NSCs *in vivo* requiring OxPhos activity, not for ATP production, but for another upstream function of the ETC. To test whether this upstream function is related to Complex I-dependent NADH oxidation, we generated transgenic *Drosophila* lines for cell-type-specific expression of LbNox, a water-forming NADH oxidase derived from *Lactobacillus brevis* ^72^ (**Figure 3C; S7E**), previously shown to also decrease the NADH/NAD^+^ balance in *Drosophila* eye discs ^65^. Ectopic expression of cytoplasmic or mitochondrial-localised LbNox partially restored NSC proliferation upon Complex I knockdown (**Figure 3F,G**), providing *in vivo* evidence for the reliance of proliferating stem cells on OxPhos primarily for NADH oxidation, rather than for ATP production.

**Figure 3.**
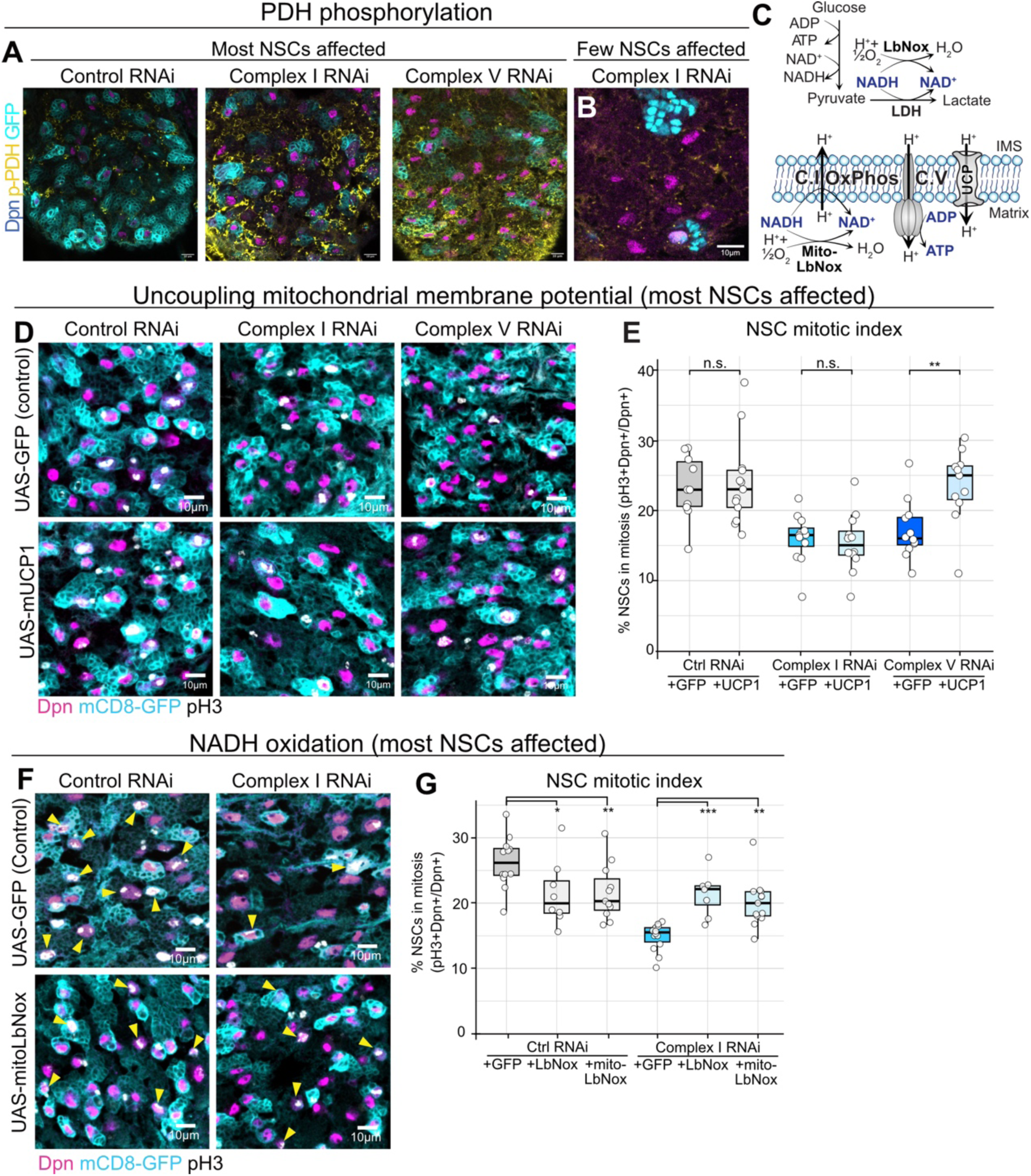
NSC proliferation relies on OxPhos to maintain redox balance. (A) Pyruvate dehydrogenase (PDH) phosphorylation (p-PDH), Dpn (NSCs) and GFP (NSC-lineages) immunostaining in L3 VNCs expressing control (UAS-*mCherry*-RNAi), Complex I (UAS-ND75 RNAi) or Complex V (UAS-ATPSynO-RNAi) RNAi in most NSCs (worniu-GAL4). (B) Pyruvate dehydrogenase phosphorylation (p-PDH), Dpn (NSCs) and GFP (NSC-lineages) immunostaining in L3 VNCs clonally expressing Complex I RNAi (LexAop-ND42 RNAi) in NSCs (Dpn-FRT-STOP-FRT-LexA). (C) Schematic of the mitochondrial electron transport chain and NADH/NAD⁺ metabolism, highlighting the effects of Complex I, Complex V, and LbNOX-mediated NADH oxidation. (D,E) Phospho-Histone H3 (pH3), Dpn (NSCs) and GFP (NSC-lineages) immunostaining (D) and mitotic index (E) in L3 VNCs expressing control (UAS-*mCherry*-RNAi), Complex I (UAS-ND75 RNAi) or Complex V (UAS-ATPSynO RNAi) RNAi in most NSCs (worniu-GAL4), while over-expressing mUCP1. Each datapoint represents an individual brain, n = 10 [Control RNAi; GFP], 13 [Control RNAi, UCP1], 12 [Complex I RNAi; GFP], 12 [Complex I RNAi; UCP1], 12 [Complex V RNAi; GFP], 11 [Complex V RNAi; UCP1]; linear mixed effect models. (F) Phospho-Histone H3 (pH3), Dpn (NSCs) and GFP (NSC-lineages) immunostaining (F) and mitotic index (G) in L3 VNCs expressing control (UAS-*mCherry*-RNAi) or Complex I (UAS-ND75 RNAi) RNAi in most NSCs (worniu-GAL4), while over-expressing Lbnox in the mitochondria (mito-LbNox) or cytoplasm (LbNox). The yellow arrows indicate the NSCs that are pH3 positive. Each datapoint represents an individual brain, n = 12 [Control RNAi; GFP], 8 [Control RNAi, LbNox], 11 [Control RNAi, mito-LbNox], 14 [Complex I RNAi; GFP], 8 [Complex I RNAi; LbNox], 13 [Complex I RNAi; mito-LbNox]; linear mixed effect models. p > 0.05 = n.s.; p < 0.05 = *; p < 0.01 = **; p < 0.001 = ***. Scale bars: 10 μm (A,B,D,F)

### Glial gap junctions enable tissue-wide non-cell-autonomous buffering

Although *Ldh* upregulation and glycolysis support ATP production to compensate for OxPhos dysfunction, the increase in NADH/NAD^+^ balance, and decrease in NSC proliferation rate when most NSCs are affected indicates that this compensation is insufficient to cell-autonomously sustain the metabolic requirements for normal proliferation. Other, likely non-cell autonomous, compensation mechanisms must therefore act to safeguard proliferation upon mosaic OxPhos dysfunction. *Drosophila* NSCs are fully enclosed in a glial niche (**Figure 4A**), consisting of three different glial subtypes: cortex glia that form a syncytial network wrapped around each NSC lineage, and perineurial and sub-perineurial glia (surface glia) that make up a highly inter-connected blood-brain barrier. ^19,79–82^ To test whether glia could provide non-cell-autonomous support for their neighbouring NSCs, we constructed a bi-partite expression system that allows clonal knockdown of OxPhos subunits in NSCs (NSC-specific flip-out LexA/LexOp ^32^), and concomitant glia-specific expression of any other RNAi via the GAL4/UAS system ^45^ (**Figure 4A,B**). We first disrupted interglial connectivity by preventing gap-junction formation with an RNAi targeting *inx2* under the control of the pan-glial driver *Repo-Gal4* (**Figure S8A**). Glial gap-junctions are required for NSC reactivation at the embryo-to-larval transition ^83^, but not for normal brain growth after reactivation ^83^. However, *inx2* knockdown in glia after NSC reactivation caused a significant non-cell-autonomous reduction in the proliferation of NSCs with clonal OxPhos dysfunction compared to those not affected by OxPhos dysfunction (**Figure 4C; S8B,C**). In addition, selective disruption of cortex glia syncytium formation ^79^ through knockdown of the known cell-cell fusion gene *sns*, but not of *WASp* or *hbs*, significantly reduced NSC proliferation with mosaic Complex I knockdown (**Figure S8D**). These results indicate that glial network integrity is required for non-cell-autonomous support of NSCs confronted with OxPhos dysfunction, possibly to allow metabolite exchange from distant glial cells with spare metabolic capacity ^16,17,84^, across a brain-wide glial network.

**Figure 4.**
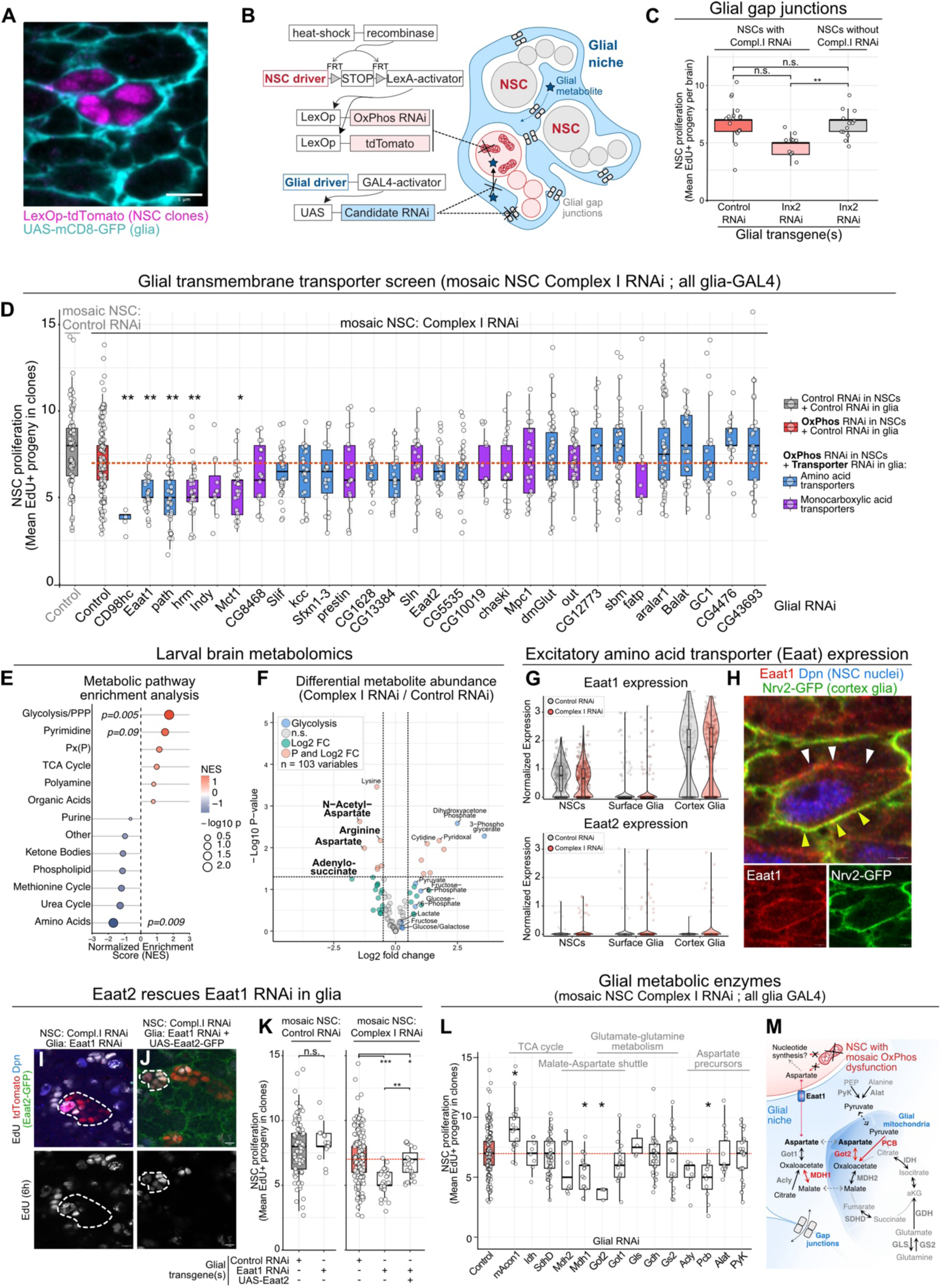
NSCs with OxPhos dysfunction rely on metabolites from a glial network. (A,B) Representative image (A) of L3 VNC with NSCs labelled with tdTomato (magenta) and the surrounding glial cells with GFP (cyan), according to the genetics described in the schematic of the bi-partite expression system (B). (C) Number of newly produced cells (EdU+, 6 hours feeding) per NSC lineage clonally expressing Complex I RNAi (LexAop-ND42 RNAi), while expressing UAS-inx2 RNAi in the surrounding glia (Repo-GAL4). Each datapoint represents number of clones within individual brains; n = 17 clones, 3 brains [Control RNAi], 10 clones, 2 brains [inx2 RNAi], 13 clones, 2 brains [inx2 RNAi; without Complex I RNAi]; linear mixed effect models. (D) NSC proliferation after knocking down transmembrane transporters in glia (Repo-GAL4) while NSCs clonally express Complex I RNAi (Dpn-FRT-STOP-FRT-LexA). NSC proliferation quantified as the number of EdU+ progeny per NSC lineage following 6hr EdU feeding. The red dashed line indicates the mean proliferation of Complex I RNAi NSCs with control RNAi expressed in glia. Each datapoint represents clones within individual brains, n = 1413 clones, 321 brains in total; linear mixed effect models. (E) Metabolic pathway enrichment analysis of differential metabolite abundance from metabolomics of whole CNS samples from L3 larvae expressing control or Complex I RNAi in most NSCs (worniu-GAL4). The x-axis shows the Normalized Enrichment Score (NES), with positive and negative values indicating enrichment toward metabolites increased or decreased in Complex I RNAi, respectively, and point size represents the nominal -log10 p-value. (F) Differential metabolite abundance between whole L3 brains expressing control or Complex I RNAi in NSCs, analyzed using limma with empirical-Bayes moderation. Each point represents a metabolite; dashed lines indicate thresholds of log2 fold change ≥ 0.5 and p < 0.05. Colours distinguish metabolites passing fold-change or *p*-value threshold, both thresholds, or neither; selected candidates labelled and glycolysis-metabolites indicated separately. (G) Normalized expression levels of *Eaat1* (top) and *Eaat2* (bottom) in NSCs, surface glia, and cortex glia under control RNAi (gray) and Complex I RNAi (red) conditions from the scRNA-seq dataset. Points represent individual cells; violin width indicates the density of the cells at each expression level; boxplots show median and interquartile range (25^th^-75^th^ percentiles) within each cell type and condition. (H) Eaat1 and Dpn (NSCs) immunostaining in L3 VNCs with cortex glia labelled by GFP (Nrv2-GFP). Yellow arrowheads indicate Eaat1 in the cortex glia membrane; white arrowhead in the NSC membrane. (I-J) EdU (6h feeding), Dpn (NSCs), tdTomato (NSC lineages) or GFP (UAS-Eaat2-GFP) immunostaining (I,J) and quantification of EdU+ progeny (6hr feeding) per NSC lineage (K) in L3 VNCs with clonal Complex I RNAi in NSCs (Dpn-FRT-STOP-FRT-LexA), and simultaneous expression of either Eaat1 RNAi alone (I) or with overexpression of Eaat2 (UAS-Eaat2-GFP) in the surrounding glia (Repo-GAL4; J). Dashed lines outline lineages from single NSCs (Dpn+, tdTomato+). Each datapoint represents number of clones within individual brains, n = 78 clones, 16 brains [mosaic NSCs: Control RNAi; Glia: Control RNAi], 16 clones, 4 brains [mosaic NSCs: Control RNAi; Glia: Eaat1 RNAi], 171 clones, 36 brains [mosaic NSCs: Complex I RNAi; Glia: Control RNAi], 28 clones, 6 brains [mosaic NSCs: Complex I RNAi; Glia: Eaat1 RNAi] and 23 clones, 6 brains [mosaic NSCs: Complex I RNAi; Glia: Eaat1 RNAi and Eaat2-GFP]; linear mixed effect models. (L) NSCs proliferation after knocking down metabolic enzymes in the glia (Repo-GAL4), while NSCs clonally express Complex I RNAi. NSC proliferation was quantified as the number of EdU+ progeny per NSC lineage following a 6h EdU feeding period. Each datapoint represents number of clones within individual brains, n = 1413 clones, 321 brains in total; linear mixed effect models. (M) Schematic illustrating the different metabolites and their contributions to metabolic pathways in glia and NSCs. Red indicates significant and grey no significant (as in L) requirement in glia to support NSC proliferation. Dashed lines indicate reactions not tested in glia. Malate-aspartate shuttle metabolites and Eaat1 shown in black. p > 0.05 = n.s.; p < 0.05 = *; p < 0.01 = **; p < 0.001 = ***. Scale bars 5 μm (A,H,I,J).

### NSCs with OxPhos dysfunction rely on glial aspartate transport

We next reasoned that membrane-transporters are likely to play a key role in a non-cell-autonomous buffering against mosaic OxPhos-dysfunction, through exchange of metabolites that can bypass auxotrophies induced by OxPhos-dysfunction ^85^. We therefore analysed glial and NSC expression of amino acid and monocarboxylate transporters in our scRNA-seq dataset (**Figure S8E,F**). Several were strongly expressed, but none showed differential expression in the glia upon NSC-specific Complex I RNAi (**Figure S5A,B; Table S2**). Taking advantage of our bi-partite expression system (**Figure 4A,B**), we then carried out an *in vivo* RNAi screen of 29 transmembrane transporters in glia, using EdU incorporation in NSC lineages with mosaic OxPhos dysfunction as a readout. Most (24/29) of these transporters were not required in glia for normal NSC proliferation (**Figure 4D**). However, glial knockdown of 5/29 transporters (*CD98hc*, *Eaat1*, *path*, *hrm* and *Mct1*) significantly decreased the proliferation rate of NSCs affected by Complex I knockdown (**Figure 4D**). Several of these glial transporters are known to be required for normal NSC proliferation in *Drosophila*, in particular the heavy chain of heterodimeric amino acid transporters, *CD98hc* ^86,87^ knockdown of which indeed strongly decreased overall brain size (**Figure S8G,H**). Glial knockdown of the alanine/glycine transporter *path* ^88^, and the aspartate/glutamate transporter Eaat1 ^89,90^ both significantly decreased proliferation of NSCs with mosaic Complex I knockdown (**Figure 4D**), but had a more subtle effect on overall brain size than glial *CD98hc* knockdown (**Figure S8G,H**). However, glial knockdown of the monocarboxylate transporters, *hrm* and *Mct1*, only decreased proliferation of NSCs with mosaic OxPhos dysfunction (**Figure 4D**), without otherwise affecting brain size (**Figure S8G,H**).

To further identify candidate metabolic pathways involved, we conducted metabolomics on whole CNS of wandering larvae expressing either Control or Complex I RNAi specifically in NSCs (**Table S10**). Pathway enrichment analysis of differential metabolite abundance revealed significant increases in glycolysis metabolites upon Complex I knockdown (p = 0.005; **Figure 4E; S8I; Table S10**), in line with our findings from the single-cell RNA-seq (**Figure 2**). High abundance of dihydroxyacetone phosphate (DHAP) and 3-phosphoglycerate metabolites, crucially involved in the glycerol-3-phosphate (G3P) shuttle ^91,92^ further support the importance of regulating NADH/NAD^+^ redox balance in NSCs upon OxPhos dysfunction. Amino acid metabolism was significantly downregulated (p = 0.009; **Figure 4E; S8I**), in particular with a strong decrease in aspartate-related metabolites, aspartate, N-acetyl-aspartate, adenylosuccinate and arginine (**Figure 4F; S8I**). A known role for OxPhos in proliferating cells is to provide sufficient NAD^+^ to support aspartate production required for nucleotide synthesis ^71,73,93,94^. One of the candidate glial transporters from our glial screen (**Figure 4D**) that therefore attracted our attention was the glutamate-aspartate transporter Eaat1, previously involved in post-synaptic glutamate recycling by astrocytes ^95^ and nutrient support for *Drosophila* NSCs during starvation ^90^. From our scRNAseq analysis (**Figure 4G**) and previous literature ^89,90^, Eaat1 is strongly expressed in NSCs and cortex glia, but not surface glia. This finding was confirmed *in vivo* by immunostaining of Eaat1 along with the NSC-marker Dpn and a genetic cortex glia marker (*Nrv2::GFP* ^96^) (**Figure 4H**). We hypothesized that exogenous supply of glial aspartate through Eaat1 might bypass the need for OxPhos-derived NAD^+^ in NSCs to support nucleotide synthesis and proliferation. *Drosophila* Eaat2 is a high-affinity aspartate and taurine transporter ^97,98^ that exhibits a stricter substrate selectivity than other known EAATs, but is not normally expressed in NSCs or their surrounding glia (**Figure 4G**) and is not required for NSC proliferation upon glial knockdown (**Figure 4D**). Using our bipartite expression system (**Figure 4B**), we found that Eaat2 overexpression could partially compensate for loss of glial Eaat1, and restore proliferation of NSCs with clonal OxPhos dysfunction (**Figure 4I-K**), indicating a key role for a metabolite that can be transported by both Eaat2 and Eaat1, including aspartate.

Finally, to understand the glial sources of aspartate synthesis, we knocked down enzymes in the glia involved in pathways related to aspartate and glutamate metabolism, and monitored proliferation of NSCs with mosaic OxPhos dysfunction (**Figure 4L,M**). None of the glial enzymes involved in the TCA-cycle (*mAcon1*, *Idh*, *SdhD*) or related to glutamate and glutamine metabolism (*Gls*, *Gdh*, *Gs2*) were required to non-cell-autonomously support proliferation of NSCs with OxPhos dysfunction (**Figure 4L**). In contrast, glial knockdown of malate-aspartate-shuttle enzymes (*Mdh1* and *Got2*) or pyruvate carboxylase (*Pcb*), involved in mitochondrial synthesis of the aspartate precursor oxaloacetate, significantly affected proliferation of NSCs with mosaic OxPhos dysfunction. Glial knockdown of several other enzymes, including ATP-citrate lyase (*Acly*), the malate-aspartate shuttle enzymes *Got1* and *Mdh2*, and the pyruvate-producing enzymes *PyK* and *Alat* did not significantly affect NSC proliferation (**Figure 4L,M**). Together, our data indicate that Eaat1-mediated aspartate transport from glia may compensate for the loss of OxPhos-dependent redox regulation in NSCs (**Figure 4M**), likely providing essential building blocks for nucleotide synthesis and proliferation.

### Glial NAD^+^ production is required to compensate for NSC OxPhos dysfunction

Synthesising the metabolites required to meet the OxPhos-induced auxotrophic demand of NSCs, including for aspartate, may impose substantial redox burden on the neighbouring glia. To test this, we perturbed glial metabolic pathways involved in redox homeostasis, and tested their impact on proliferation of NSCs with mosaic OxPhos dysfunction. Glial knockdown of OxPhos Complex I or V subunits did not affect proliferation of wild-type NSCs (**Figure S9A**) but did prevent NSCs with clonal OxPhos dysfunction from proliferating normally (**Figure S9A**). In contrast, glial RNAi against *DGAT1, Lsd2* or *Pld*, to inhibit glial lipid droplet formation, previously shown to protect NSCs from environmental oxidative damage ^99^, did not (*DGAT1*, *Lsd2*) or only slightly (*Pld*) impact proliferation of NSCs with clonal OxPhos dysfunction (**Figure S9A**). We next tested the role of LDH, which is strongly expressed in most glial cell-types, in particular surface glia (**Figure 2I**). Just like for glial OxPhos subunits (**Figure S9A**), glial *Ldh* knockdown did not affect proliferation of wild-type NSCs (**Figure 5A-E**), but did significantly decrease proliferation of NSCs with clonal OxPhos dysfunction (**Figure 5B,D,E**). Glial LDH and OxPhos are thus not normally required for NSC proliferation, but become essential to support NSC proliferation upon OxPhos inhibition, indicating that the metabolic demands of NSCs with OxPhos dysfunction non-cell-autonomously deplete the spare metabolic capacity of glia.

**Figure 5.**
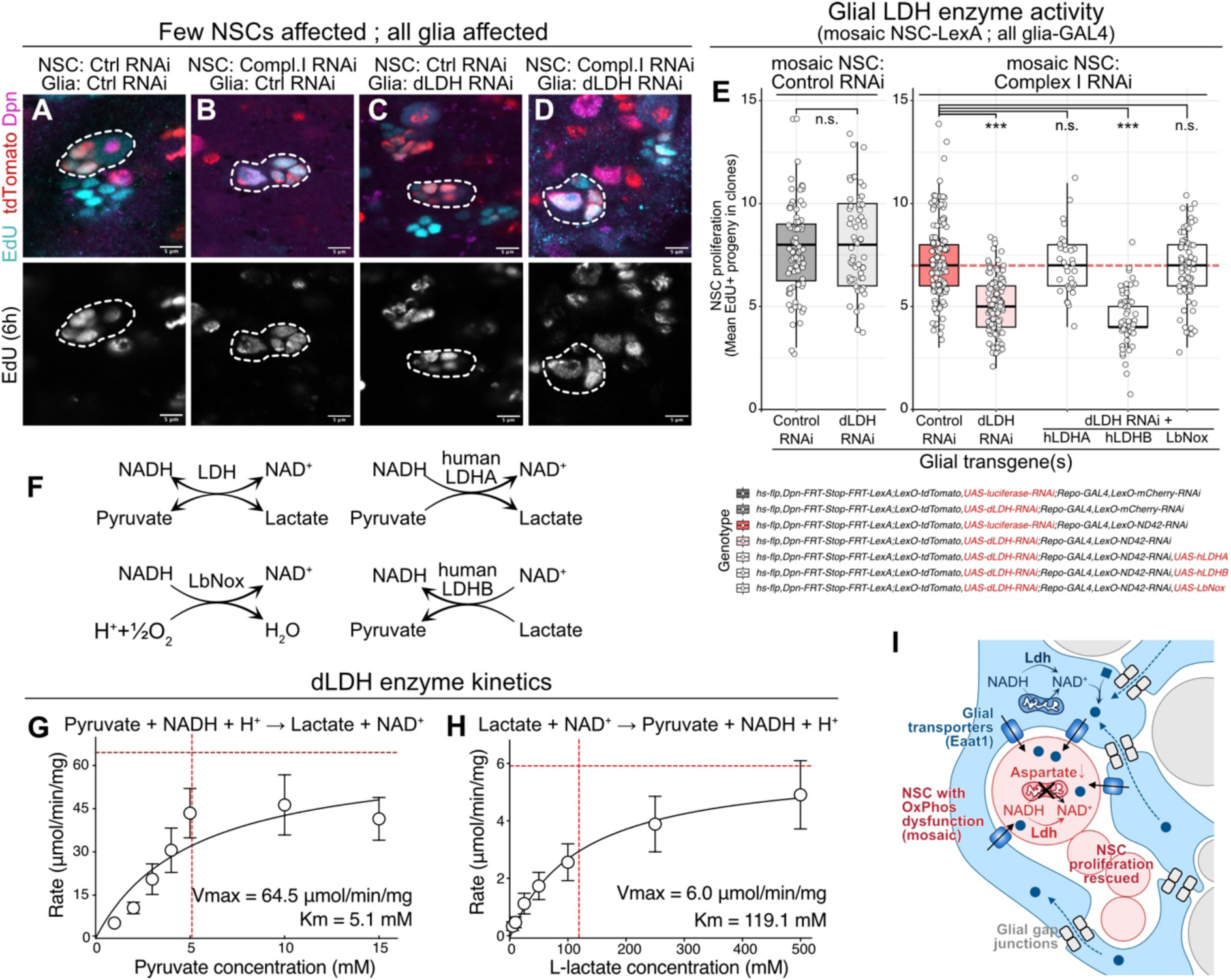
Glial NAD^+^ production is required to compensate for NSC OxPhos dysfunction. (A-D) EdU (6hr feeding, NSC progeny), Dpn (NSCs) and tdTomato (NSC lineages) immunostaining in L3 VNCs with clonal Complex I RNAi (LexAop-ND42 RNAi) in NSCs (Dpn-FRT-STOP-FRT-LexA), and control (UAS-Luciferase RNAi; A,B) or Ldh RNAi (C,D) in surrounding glia (Repo-GAL4). Dashed lines outline lineages from single NSCs. (E) NSC proliferation (number of newly produced cells (EdU+, 6hr feeding) per NSC lineage expressing the indicated genotypes. Each datapoint represents number of clones within individual brains, n = 78 clones, 16 brains [mosaic NSCs: Control RNAi; Glia: Control RNAi], 58 clones, 12 brains [mosaic NSCs: Control RNAi; Glia: LDH RNAi], 171 clones, 36 brains [mosaic NSCs: Complex I RNAi; Glia: Control RNAi], 114 clones, 21 brains [mosaic NSCs: Complex I RNAi; Glia: LDH RNAi], 34 clones, 7 brains [mosaic NSCs: Complex I RNAi; Glia: LDH RNAi and hLdhA], 53 clones, 12 brains [mosaic NSCs: Complex I RNAi; Glia: LDH RNAi and hLdhB], 73 clones, 18 brains [mosaic NSCs: Complex I RNAi; Glia: LDH RNAi and LbNox]; linear mixed effect models. (F) Schematic representation of the LDH reaction, illustrating LDH in general, human LDHA, human LDHB, and LbNox, illustrating their respective interactions with the NADH/NAD⁺ redox couple. Arrows indicate directionality of the corresponding redox reactions. (G,H) NADH absorbance assay using different substrate concentrations and Michaelis-Menten kinetics applied to determine the Vmax and Km, carried out for pyruvate (G) or lactate (H) at pH7.4. Error bars: SEM. (1) Schematic illustrating how glial gap junction-mediated metabolite exchange, glial transmembrane transporter (e.g. Eaat1) mediated metabolite transfer to NSCs, and Ldh upregulation in NSCs, may contribute to NSC redox balance and proliferation, when confronted with OxPhos dysfunction and aspartate depletion. p > 0.05 = n.s.; p < 0.05 = *; p < 0.01 = **; p < 0.001 = ***. Scale bars: 5 μm (A-D).

LDH can either oxidise lactate or reduce pyruvate, producing NADH or NAD^+^, respectively (**Figure 5F**). In many species, including in all vertebrates, different LDH isoforms have different substrate preferences and reaction kinetics ^100,101^, but *Drosophila* has only one isoform, which is presumed to catalyse both reactions ^7,21,91^. We initially reasoned that glial LDH might be involved in removing excess lactate secreted by NSCs upon OxPhos dysfunction, in a mechanism reminiscent, but opposite to the neuron-glia lactate shuttle ^20,21,102,103^. However, when we measured *in vitro* recombinant LDH enzyme activity with pyruvate or lactate as substrates, *Drosophila* LDH was found to have a >20 x higher affinity for pyruvate (Km 5.1 mM, 95%CI 2.5-10.9; Vmax 64.5 μmol/min/mg, 95%CI 47.8-93.4) than for L-lactate (Km 119.1 mM, 95%CI 67.1-219.9; Vmax 6.0 μmol/min/mg, 95%CI 4.9-7.7) (**Figure 5G,H; S9B,C**). The Km for pyruvate of *Drosophila* LDH was higher than of human LDHA ^104^, but in line with a previous report ^105^, indicating that *Drosophila* LDH has a strong substrate selectivity for pyruvate over lactate. We next obtained *Drosophila* strains that allow ectopic expression of the human LDH isoforms LDHA or LDHB (**Figure 5F**). which are thought to preferentially reduce pyruvate or oxidise lactate in a physiological environment ^106,107^. We expressed either isoform, or the NADH oxidase LbNox ^72^ (**Figure 5F**), in glia, while knocking down endogenous glial *Drosophila Ldh* expression, and measured proliferation of NSCs with mosaic Complex I RNAi. To our surprise, LDHA and LbNox, but not LDHB, could rescue the non-cell-autonomous effect of glial *Ldh* knockdown, and restore NSC proliferation (**Figure 5E; S9D-F**). We also confirmed that ectopic expression of Lbnox in the glia, in a Complex I RNAi background in the NSCs, doesn’t have an effect in the NSCs proliferation (**Figure S9G-I**). This means that, in response to NSC OxPhos dysfunction, glial dLDH most likely does not consume but instead produces lactate. Moreover, because glial LbNox-expression can bypass the requirement for glial LDH to support proliferation of NSCs with mosaic OxPhos dysfunction, it is the resulting glial NAD^+^ that is required to non-cell-autonomously restore NSC proliferation (**Figure 5I**).

## Discussion

Cells in a heterogeneous tissue-environment cooperate by dividing metabolic pathways between cell-types. In the CNS, for example, glial cells are thought to convert glucose to lactate, which fuels OxPhos in neighbouring neurons ^20–22,106^. However, it remains unclear whether this metabolic compartmentalisation can adapt dynamically to perturbations, and whether it creates tissue-wide spare metabolic capacity that could underlie organ-specific vulnerability or resilience to environmental fluctuations, stress or disease.

Here, we identified a tissue-wide non-cell-autonomous metabolic buffering mechanism that allows NSCs in the developing *Drosophila* brain to sustain proliferation and normal developmental patterning despite mitochondrial dysfunction. While NSCs compensate for OxPhos deficiency by activating glycolysis and LDH-mediated lactate and NAD^+^ production, we find that this is insufficient to cell-autonomously sustain normal proliferation rates *in vivo*.

Instead, OxPhos dysfunction, and the auxotrophies it creates, render NSCs dependent on non-cell-autonomous supply of exogenous metabolites from surrounding glial cells. Through a small-scale RNAi-screen, we discovered five glial transmembrane transporters required for bypassing the NSC redox imbalance, including the excitatory amino acid transporter, Eaat1, potentially involved in aspartate exchange for NSC nucleotide synthesis. Crucially, this non-cell-autonomous compensation mechanism relies on glial OxPhos- and LDH-mediated NAD^+^ production, which become exhausted when too many NSCs across the brain are affected. OxPhos dysfunction thus creates a redox-limited biosynthetic state in NSCs that neighbouring glia can bypass through metabolic cooperation.

We propose a model whereby a brain-wide network of different glial cell types that make up the stem cell niche, connected through cell fusion and glial gap junctions, contains sufficient spare metabolic capacity to bypass OxPhos requirement in a few NSCs (**Figure 5I**). Cortex glia immediately surrounding a dysfunctional NSC can generate excess aspartate by cooperating with distant brain cells and ‘crowd-sourcing’ sufficient redox metabolites, similar to what was shown for gap-junction-mediated nucleotide sharing in the wing disc ^84^ or for metabolite exchange in various forms of human cancer ^108,109^. However, this extended support network progressively decompensates when there are too many ‘mouths to feed’, leading to NSC cell cycle slowing and developmental defects ^37,38,42^.

A naturally occurring, and ubiquitous form of mitochondrial mosaicism is caused by heteroplasmic mtDNA variation ^110,111^. Taking advantage of a heteroplasmic *Drosophila* strain affecting Complex IV activity ^47^ and our recently-developed mtDNA-smFISH ^48^ to quantify *in situ* heteroplasmy levels, we observed similar compensation mechanisms to act in heteroplasmic brains: NSCs with high levels of mutant mtDNA can sustain normal proliferation, but only when surrounded by sufficient cells with lower mutation load. In mitochondrial disease, and during aging, mtDNA mutations accumulate in specific tissues, to cause disease when the mutation load within each cell reaches a certain biochemical threshold ^112^. Our data now suggest that this threshold is not simply cell-type specific, but can be modulated by each cell’s direct tissue-environment. This may explain the finding of cells with near-homoplasmic pathogenic mutation levels in recent single-cell sequencing studies ^27,28^. Further tissue-specific resilience to mitochondrial disease is likely to arise from cell-type composition, in particular in the brain, where the glia/neuron-ratio in the cerebral cortex is ∼15 times higher than in the cerebellum. ^113^ The response to mitochondrial dysfunction differs between neurons and glia ^114–116^, and glia have been shown to play a protective role against neuronal oxidative stress ^117–119^. Differences in cell-type-composition and spare metabolic capacity may thus render specific brain-regions more, or less, vulnerable to mitochondrial disease, similar to what was previously suggested for Huntington’s disease ^120^. Recent progress in combined RNA- and mtDNA single-cell sequencing will help address these questions, by measuring and perturbing the stress response to mtDNA mutations in different tissue-contexts.^26,31,110^

How NSCs signal their metabolic requirements to surrounding glia, and then trigger redistribution of metabolites throughout the glial network, remains unknown. Within NSCs, we identified a range of transcription factors, including the *Drosophila* ATF4-homolog *crc* ^60^ and several NSC-specific factors like *Ets65a* and *E(spl)m3* ^50^, that regulate a cell-autonomous stress-response and *Ldh* gene expression in response to Complex I knockdown. However, single-cell RNA-seq did not reveal major changes in glial gene expression upon NSC-specific OxPhos dysfunction, also not for the transporters and enzymes that we found to be required for non-cell-autonomous buffering. Nevertheless, mitochondrial dysfunction in one cell is well known to affect the behaviour of surrounding cells in many different contexts, such as tumourigenesis or life-span regulation ^121–124^. Whether similar signals act in the *Drosophila* brain, and whether these can be exploited for therapeutic interventions to promote resilience to disease remains to be explored.

Our data provide *in vivo* genetic evidence for the importance of OxPhos to maintain the NAD^+^/NADH redox balance in proliferating cells. OxPhos dysfunction likely creates auxotrophy for aspartate to support nucleotide synthesis, similar to previous observations in cell culture and tumour models ^10,70,71,73,93,94^. This is evidenced by preferential depletion of aspartate-related metabolites in metabolomics of brains where most NSCs are affected by OxPhos dysfunction, and the requirement for glial expression of malate-aspartate shuttle enzymes and the glutamate-aspartate transporter Eaat1. To our surprise, when confronted with OxPhos dysfunction in neighbouring NSCs, glia also became reliant on LDH. However, this was not to revert NSC-derived glycolytic lactate back to pyruvate, akin to a reverse lactate shuttle ^125^. Instead, we found that the single *Drosophila* LDH isoform most likely only reduces pyruvate, and it is the resulting NAD^+^ that is required to support NSC proliferation, presumably by enabling excess glial aspartate production. It will be interesting to study how LDH with unidirectional activity nevertheless enables a neuron-glia lactate shuttle in *Drosophila* ^21,22,126–128^, whether certain modifications or circumstances modulate or invert *Drosophila* LDH activity, or whether another, as-yet unidentified dehydrogenase might consume lactate in *Drosophila*, similar to the ability of LDH isoforms and other dehydrogenases to also convert 2-oxo-glutarate into L-2-hydroxyglutarate in hypoxic conditions ^105,129,130^.

Our findings indicate that during development, heterogeneity within a tissue ensures robustness in outcome, by providing a buffer against stochasticity and perturbations from signalling, metabolic and environmental variation. The mechanisms and principles we describe in the developing *Drosophila* brain are likely relevant in many other contexts, ranging from heterogeneous bacterial communities ^13–15^ to human cancer ^10,12^. Our findings suggest that vulnerability to perturbations is not solely encoded within individual cells, but emerges from the metabolic architecture of the tissues they inhabit. It underscores the importance of diversity across biological systems, and the systemic robustness created through equal redistribution of spare capacity. Exploring these mechanisms to enhance resilience to disease may offer promising future therapeutic targets, including for rare mitochondrial diseases that remain so far without cure.

## Acknowledgements

We thank all lab members and P. Speder, H. Ma, J. Prudent, D. Atwell, W. Dunn, E. Contreras and W. Staels for helpful discussions and advice; P.F. Chinnery, R. Horvath and H. Biggs for continuous support and interest; the Cambridge Fly Genetics Facility and FlyORF for *Drosophila* injection services; S.R. Chowdhury from the MRC MBU Imaging Facility for support with imaging; R. Schulte, G. Grondys-Kotarba from the CIMR Flow Cytometry Facility and A. Glynos for assistance with cell sorting; the University of Birmingham Metabolic Tracer Analysis Core for support and resources; J. Knoblich, D. van Meyel, P. MacDonald for antibodies; T. Lee, M. Landgraf, T.D. Southall, I. Lohmann, D. van Meyel, H. Ma, C. Collins, Y-W. Fridell for transgenic *Drosophila* lines; V. Mootha for LbNox plasmids (Addgene 75285 and 74448).*Drosophila* stocks obtained from the Bloomington *Drosophila* Stock Center (NIH P40OD018537), Kyoto Drosophila Genomics and Genetics Resources (DGGR) and Vienna *Drosophila* Resource Center (VDRC) were used in this study.

## Funding

Wellcome Clinical Research Career Development Fellowship 219615/Z/19/Z (JvdA)

Evelyn Trust Medical Research Grant 21-25 (JvdA)

UKRI BBSRC Responsive Mode Research Grant BB/X00256X/1 (JvdA)

Wellcome Discovery Award 226653/Z/22/Z (JvdA)

Medical Research Council National Mouse Genetics Network MC_PC_21046 (JvdA)

Medical Research Council Mitochondrial Biology Unit MC_UU_00028/8 and MC_UU_00028/2 (JvdA, ERSK)

Rosetrees Trust PGL23/100048 (JvdA)

LifeArc 10478 (JvdA)

Muscular Dystrophy UK 23SI-PRG60-0013 (JvdA)

University of Cambridge DTP in Medical Research and Cambridge Trust studentship (TE)

Tenure Track Fellowship from the University of Liverpool (MS)

For the purpose of open access, the authors have applied a Creative Commons Attribution (CC BY) license to any Author Accepted Manuscript version arising from this submission.

## Author contributions

Conceptualization: JvdA

Methodology: SP, DD, RC, HD, AHA, MS, JvdA

Investigation: SP, DD, RC, HD, NBM, BAW, TE, BM, AHA, SAJ, MSK, JvdA

Formal analysis: SP, AD, DD, MSK, JvdA

Visualization: SP, AD, DD, HD, MSK, JvdA

Software: RB

Supervision: MS, ERSK, DAT, JvdA

Funding acquisition: JvdA

Writing – original draft: SP, AD, JvdA

Writing – review & editing: all authors

## Data, code and materials availability

### Lead contact

Requests for further information and resources should be directed to and will be fulfilled by the lead contact, Jelle van den Ameele

## Materials availability

All materials, resources and reagents are listed in the methods section and Table S9. Further requests should be directed to and will be fulfilled by the lead contact. **Data and code availability**: Single-cell sequencing data will be made publicly available upon publication on GEO, with analysed data available in Tables S1-8. Metabolomics results are available in Table S10. Code for single-cell sequencing and image analysis is available on Github (https://github.com/JvdAlab/Drosophila-NSC-mosaicism).

## Materials and Methods

### Fly husbandry

*Drosophila melanogaster* strains were maintained at 25 °C. For most experiments, embryos were collected on cornmeal plates for 4 hr and kept at 25 °C until analysis. Unless indicated otherwise, larvae were matched for developmental timing at wandering third instar (L3). For time-course experiments, embryos were collected on cornmeal plates and larvae were transferred to a fresh food vial within 2 hr of hatching (designated 0 hr ALH) and grown at 25°C until the desired stage. For clonal analysis, embryos and larvae were grown at 25 °C and heat-shocked when indicated for 20 min in a 37 °C water bath.

### Fly stocks

UAS-mCherry-TRIP (Bl#35785) was used as control RNAi throughout the study. Unless otherwise indicated, all UAS-Complex I RNAi data are from ND75-TRIP (NDUFS1; Bl#33911; TRiP.HMS00854) ^124^ and all Complex V RNAi data from ATPsynO-TRIP (Bl#43265; TRiP.GLC01454). For the metabolomics experiment (**Figure 4E,F; S8I**) UAS-ND42-TRIP (Bl#32998; TRiP.HMS00798) was used instead. All LexAOp-RNAi lines against OxPhos components were generated in this study based on TRiP RNAi sequences ^131^: LexAOp-mCherry-RNAi, LexAOp-ND42-RNAi (based on TRiP.HMS00798) and LexAOp-ATPSynO-RNAi (based on TRiP.GLC01454). The NSC GAL4-driver used throughout the study was Worniu-GAL4 on II ^44^ (Bl#56553), either on its own, or recombined with UAS-mCD8-GFP on II. The genotypes for the generation of flip-out clones for the expression of Complex I RNAi in a few NSCs (**Figure 1C; S1H**) were as follows: y,w,hsflp;+;UAS-ND75-RNAi crossed to w;UAS-mCD8GFP;Actin-FRT-STOP-FRT-GAL4 (AyGAL4). The genotypes for the expression of Complex I RNAi in all NSCs within the brain (**Figure 1A, S1G**) were as follows: y,w,hsflp;+;UAS-ND75-RNAi crossed to w;worniu-GAL4;Tub-FRT-STOP-FRT-GFP, which randomly labelled GFP clones while expressing the RNAi in all the NSCs. The genotypes for the generation of flip-out clones for the screening experiments (**Figure 4D,I,J,K,L; S8D,G,H; 5A-D, E; S9A,D-F, G-I**) were as follows: y,w,hsflp;+;UAS-Screening RNAi crossed to Dpn-FRT-STOP-FRT-LexAp65;LexAOp-NLS-tdTomato;Repo-GAL4,LexAOp-ND42-RNAi for Complex I knock down and Dpn-FRT-STOP-FRT-LexA;LexAOp-NLS-tdTomato;Repo-GAL4,LexAOp-mCherry-RNAi as a control. This resulted in NSC lineages that were randomly marked upon heat-shock, knocking down the Complex I subunit ND-42, whereby all the surrounding glial cells expressed the desired RNAi. Sources from these transgenes are: y,w,hsflp (Bl#26902), Dpn-FRT-STOP-FRT-LexA (Bl#56162), LexAOp-NLS-tdTomato (Bl#66690 or Bl#66691), Dpn-LexAp65 (gift from Tzumin Lee) ^32^, UAS-mCD8GFP, AyGal4 (Bl#4411), Repo-GAL4 (Bl#7415), Tub-FRT-Stop-FRT-Gal80 (Bl#38880).

All UAS-RNAi lines used in this study, and the results obtained from the glial RNAi-screen (**Figure 4D, K, L; S8D; S9A**) are summarised in **Table S9**. For the innexin experiment (**Figure 4C; S8B-C**) the genotypes were as follows: y,w,hsflp;Tub-GAL80^ts^;UAS-inx2-RNAi crossed to Dpn-FRT-STOP-FRT-LexA;LexAOp-NLS-tdTomato;Repo-GAL4,LexAOp-ND42-RNAi; embryos and larvae were grown at 25°C and heat-shocked after NSC reactivation at 48h ALH for 20 min in a 37°C water bath followed by incubation at 29°C. Heteroplasmic *Drosophila*, carrying a combination of wild-type *D. yakuba* mtDNA and mutant *D. melanogaster* mtDNA (mt:ND2^del1^+mt:COI^T00I^), on a nuclear *D. melanogaster* w1118 genomic background (gift from Hansong Ma) were described previously ^47,132^. To make clones of mtDNA depletion (Figure 1L-P), the Tub-FRT-CD2-FRT-Gal4,UAS-GFP line (FBtp0010169) was used. For the scRNA-seq experiment (**Figure 2A**), worniu-GAL4 (Bl#56553) was combined with UAS-mCD8-GFP (Bl#5137) to label NSCs, while Repo-LexA::GAD driver (FBtp0041066) was combined with LexAop-tdTomato (Bl#66691) to label glial cells. Other lines used throughout the study: UAS-hLdhA (Bl#79190), UAS-hLdhB (Bl#79193), UAS-dLdh (FlyORF#F002924), Ldh::GFP (gift from Ingrid Lohmann) ^55,60^, UAS-mUCP1 (gift from Catherine Collins and Yih-Woei Fridell) ^133^, UAS-hUCP2 (This study), UAS-SOD2 (Bl#24494), UAS-Eaat2 (gift from Donald van Meyel) ^134^, Nrv2::GFP (Bl#6828), NP2222-GAL4 (DGGR#112830), UAS-LbNox (This study), UAS-traNLS-LbNox (This study), UAS-mito-LbNox (This study).

### Plasmids

UAS-LbNox (cytoplasmic localisation), UAS-2xSV40NLS-LbNox (cytoplasmic and weak nuclear localisation), mito-LbNox (mitochondrial localisation, human COXIV mitochondrial targeting signal), UAS-TraNLS-LbNox (nuclear localisation) were cloned by PCR from Addgene plasmids 75285 or 74448 ^72^, including a C-terminal Flag-tag, and inserted into NotI-XbaI of pUAST-attB from DGRC_1419 ^135^. Nuclear localisation signals were inserted N-terminal; TraNLS ^136^ (5’-aagagcaggcacagaaggcatcgccagcgctctaggagccgcaatcgcagccgaagtcgcagcagtgaacgaaaacgccgtcaacggagccgaagtcgcagcagtgaacgaagacgc-3’) was amplified from *D. melanogaster* gDNA using primers listed in **Table S11**, and inserted into NotI-EcoRI of pUAST-attB-LbNox.

dLDH for recombinant expression: Sequences for lactate dehydrogenase from *Drosophila* (uniprot accession code: Q95028) with an N-terminal strep-tag (WSHPQFEK) were synthesised by GenScript in the pET-21a(+) expression vector to obtain pET-21a-dLDH plasmids. Human UCP2 was amplified from HeLa cell cDNA using primers listed in **Table S11** and cloned into NotI-NdeI of pUAST-attB.

shRNA hairpins targeting *mCherry*, *ND-42*, or *ATPsynO* were designed based on the corresponding TRiP RNAi sequences ^131^: mCherry (Bl#35785), ND42-RNAi (BDSC #32998; TRiP.HMS00798) and ATPsynO-RNAi (BDSC #43265; TRiP.GLC01454) and annealed oligonucleotides encoding the hairpin with NheI and EcoRI overhangs were ligated into NheI-EcoRI of pWALIUM20 (DGRC_1475) downstream of the LexAop promoter. Correct insertion and orientation were confirmed by sequencing. Plasmid DNA was obtained using QIAGEN Plasmid Plus Midi Kit (QIAGEN CAT#12945) and sent for microinjection (Cambridge Fly Facility) into the y,v,nos-int;;attP2 (used in this study) or y,v,nos-int;attP40 strains.

### Immunostaining, mtDNA-smFISH-HCR, EdU

Larval brains were dissected in PBS followed by fixation with 4% PFA (formaldehyde/PBST (PBS with 0.3% Triton)) for 20 min and washed three times in PBST in an orbital shaker. For EdU incorporation, L3 larvae were placed in instant food containing EdU where they were left feeding for 6 hr. After that, the larval brains were dissected followed by fixation as usual. After incubation with primary and secondary antibodies, the brains were incubated with the EdU reaction cocktail for 30min at room temperature (1X Click iT Reaction buffer, CuSO4, Alexa Azide 647, 1X Reaction buffer additive). The following primary antibodies were used: rat anti-PH3 (1/1000, Abcam ab10543); chicken anti-GFP (1/500, Abcam ab13970); rabbit anti-RFP (1/500, Abcam ab62341); rat anti-Dpn (1/100, Abcam ab195172); guinea pig anti-Dpn (1/1000, gift from Jurgen Knoblich ^137^); mouse anti-ATP5B (1/500, Abcam ab14730), rabbit anti-Eaat1 (1/5000, gift from Donald van Meyel ^138^); rabbit anti-pH3 (1:800, Thermo Ficher Scientific PA5-17869); rabbit p-Pdh (1/500, Cell Signalling 37115), rabbit p-AMPK (Cell Signalling Technology 2531), rabbit anti-FLAG (Cell Signalling D6W5B), rabbit anti-Imp (gift from Paul MacDonald ^139^), rabbit anti-LDHA (1/500, Boster Bio DZ41222). Secondary antibodies for conventional immunostaining were Alexa Fluor 405, 488, 546, 647 -conjugated secondary antibodies raised in goat or donkey (1/500, Life Technologies). Single-molecule mtDNA fluorescent in situ hybridisation with hybrdisation chain reaction v3.0 (mtDNA-smFISH-HCR) was performed according to the protocol as described previously ^48^. Tissues were mounted in ProLong Diamond antifade mounting medium (Thermo Fisher Scientific P36961). Slides were stored at 4℃ and imaged after 24 hours.

### Dihydroethidium (DHE) and Tetra-methylene-rhodamine-methyl-ester (TMRM) Imaging

For DHE treatment, third instar larval brains were dissected in 1X phosphate-buffered saline (PBS). Dissected brains were then incubated in 1X PBS containing 30 µM DHE for 15 minutes at room temperature, protected from light. Following incubation, samples were washed briefly in 1X PBS and mounted on the Lab-Tek chamber containing Schneider’s medium. Brains were immediately subjected to live imaging. Live imaging of DHE and mCD8-GFP fluorescence was performed using a Zeiss LSM 880 confocal microscope with 63x objective. DHE fluorescence was detected using the RFP filter, while mCD8-GFP was visualized using the EGFP filter. Imaging conditions such as laser power, acquisition speed, frame size and gain were kept constant across all samples.

For TMRM treatment, third instar larval brains were dissected in Schneider’s medium and incubated with TMRM (100 nM; Thermo Fisher Scientific) for 30 minutes at room temperature. Following incubation, brains were mounted on a Lab-Tek chamber containing Schneider’s medium and imaged immediately. Live imaging was performed using a Zeiss LSM 880 confocal microscope equipped with a 63× oil immersion objective. TMRM fluorescence was detected using the RFP filter. Imaging parameters were kept identical across all genotypes.

### Recombinant LDH expression, purification and validation

pET-21a-LDH constructs were transformed into the *Escherichia coli* strain BL21(DE) by electroporation using standard methods. Transformed bacteria were cultured in 1 L terrific broth (supplemented with 100 µg/mL ampicillin, 0.1% glucose, 1 mM MgCl_2_, 1.0% glycerol) at 37 °C until OD_600_ = 0.7. Cells were induced with 1 mM IPTG (Thermo Scientific) for 4 hours, harvested by centrifugation (4,000g, 20 min, 4 °C) and resuspended in lysis buffer (150 mM NaCl, 50 mM Tris-HCl pH 8, 5 mM MgCl_2_, 10% glycerol, 10 µg/mL DNAse I and a Complete Mini EDTA-free protease inhibitor tablet (Roche)). The bacteria were disrupted mechanically with a cell disruptor (Constant Cell Disruption Systems) at 33 kpsi and centrifuged (205,000g, 30 min, 4 °C). The supernatant was collected and incubated with 2 mL Streptactin XT 4Flow high-capacity slurry (IBA) for 1 hour at room temperature. The resin was washed with 50 mM Tris-HCl pH 8.0, 150 mM NaCl, and protein eluted in a buffer containing 50 mM Tris-HCl pH 8.0, 150 mM NaCl, 50 mM biotin. The protein was desalted into 50 mM Tris-HCl pH 8.0, 150 mM NaCl using a PD-10 column (Cytiva), and the protein concentration determined by nanodrop (extinction coefficient = 39.2 M^-1^ cm^-1^; molecular mass = 36.8 kDa) and BCA. Differential scanning fluorimetry (nanoDSF) was used to assess the thermostability of the protein. For the analysis, 10 μg of protein (which contains six tryptophans) was loaded into NanoDSF-grade glass capillaries and analysed on a Prometheus NT.48. The temperature was progressively increased from 25 °C to 95 °C at a rate of 4 °C per minute, and the apparent melting temperature (Tm) determined by the PR.ThermControl software (NanoTemper Technologies). The monomer weight was validated by SDS-PAGE. To confirm the protein formed tetramers, analysis by size exclusion chromatography (SEC) was performed. In brief, 150 μL of purified LDH was applied to a Superdex 200 Increase 10/300 SEC column (GE Healthcare) equilibrated with purification buffer on an ÄKTA explorer system (Amersham). The determination of the apparent molecular weight of LDH was done by comparing the elution volume to those of protein standards of known molecular mass in gel filtration calibration kits (GE Healthcare).

### LDH activity assay

A SpectraMax ABS Plus was used for cuvettes and a CLARIOstar Plus used for measuring activity in 96-well plates. NADH oxidation (the pyruvate to lactate reaction) or NAD+ reduction (the lactate to pyruvate reaction) was monitored at 340–380 nm (e = 4.81 mM^-1^ cm^-1^). A 2X mix of LDH and pyruvate or lactate was added to the plate before adding an equivalent volume of 2X mix of NADH or NAD^+^ to initiate the reaction. Reactions were performed in 200 mM Tris-HCl pH 7.4. Pyruvate assays used 776 μM NADH and 2 μg/mL dLDH. Lactate assays used 7.76 mM NAD^+^ and 10 μg/mL dLDH. All measurements were performed at 30 °C. The absorbance was measured every 15 seconds, and the initial rate calculated using linear regression from the first 20 measurements (five minutes) according to the Beer-Lambert Law, after subtracting background absorbance. Kinetic parameters were determined in Prism (GraphPad) by modelling the experimental data using the Michaelis Menten function.

### Metabolomics and analysis

For sample preparation, 10mg brains was collected from third instar larvae, with six biological replicates per condition. Brains were dissected in ice cold PBS followed by centrifugation at 300 g for 3 min at 4°C. Supernatant was carefully discarded, samples were stored on dry ice and were sent for LC-MS processing. Metabolomics was conducted by liquid chromatography-mass spectrometry (LC-MS) as described previously. ^140^ An extraction solvent comprising 80:20 (v/v) LC-MS grade methanol (LiChrosolv) and LC-MS grade water (Milli-Q) was prepared and supplemented with 0.25 μg/mL D6-glutaric acid as an internal standard. The solution was stored on ice prior to use. Pre-chilled extraction solution (250 μL) was added to each extraction tube containing homogenisation beads and sample material. Samples were homogenised using a TissueLyser III (Thermo Fisher Scientific) at 30 Hz for 60 s, employing a pre-chilled (-80°C) tube holder and mount. Following homogenisation, lysates were centrifuged (20,000*g, 5 min, 4°C) to pellet insoluble cellular debris and proteins. A 200 uL aliquot of the supernatant was carefully transferred to a 2 mL wide-bottom microcentrifuge tube, ensuring the pellet remained undisturbed and maintaining precise volume consistency across samples. Supernatants were evaporated to dryness at room temperature in a SpeedVac vacuum concentrator (Thermo Fisher Scientific). The remaining protein pellets were air-dried overnight in a fume hood and quantified the following day using a bicinchoninic acid (BCA) assay to enable LC-MS data normalisation.

Derivatisation reagents were prepared fresh in a fume hood prior to use: 3-nitrophenylhydrazine (3-NPH; 33 mg/mL in 75:25 v/v methanol/water), 1-ethyl-3-(3-dimethylaminopropyl) carbodiimide (EDC; 20 mg/mL in 100% methanol), and pyridine (2.5% v/v in 100% methanol). Dried metabolite residues were reconstituted in 50 uL of LC-MS grade water, followed by the sequential addition of 50 μL each of the 3-NPH, EDC, and pyridine solutions. Tubes were tightly capped, vortexed for a minimum of 15 s to ensure complete resuspension and incubated on ice for 1 h protected from light. Following derivatisation, samples were dried to completeness at room temperature in a SpeedVac system. The dried derivatised residues were resuspended in 50 μL of 25:75 (v/v) methanol/water with vigorous vortexing and transferred to LC-MS vials for analysis.

LC-MS analysis was performed using a Waters Acuity Premier HPLC system coupled to a Waters Xevo TQ-XS. Analyte separation was achieved using a Waters Premier C18 column (1.7 µm, 2.1 x 150 mm) with 0.01% formic acid (LiChropur) in LC-MS grade water (mobile phase A) and LC-MS grade acetonitrile (Biosolve) with 0.01% formic acid (mobile phase B). The LC flow was set at 0.35 mL/min and the gradient was held at 99% A for 5 minutes, 94% A at 6 minutes and held for 10 minutes, 90% A at 22 minutes, 85% A at 25 minutes, 75% A at 30 minutes, 70% A at 35 minutes, 0% A at 40 minutes and held for 3 minutes, then returning to 99% A at 43.5 minutes and stabilising for 1.5 minutes.

Data were post-processed using Skyline (MacCoss Lab). Data normalisation (to internal standard and protein abundance) was performed using an in-house R Shiny app. Metabolomics differential abundance analysis was performed using metabolite abundances normalized to the internal standard followed by total-sum normalization. Abundance values were log2-transformed with a small offset, and differential abundance between ND-42 KD and control samples was tested using limma v3.58.1. A linear model was fit for each metabolite and ND-42 KD abundance was contrasted to the control abundance with empirical Bayes moderation applied. Differentially abundant metabolites were identified using a threshold of p < 0.05 and absolute log2 fold change ≥ 0.5. Pathway-level significance was assessed by gene set enrichment analysis (fgsea v1.32.4), using a signed -log10(p-value) ranking metric, with pathway annotations defined by metabolite class.

### Single-cell sequencing

For sample preparation, worniu-GAL4 was combined with UAS-GFP to label neural stem cells, while Repo-LexA was combined with LexAop-tdTomato to label glial cells. These flies were then crossed to either UAS-mCherry-RNAi (BDSC #35785) as the control or UAS-ND75-RNAi (BDSC #33911; TRiP.HMS00854) to induce ND-75 knockdown. Second instar larvae (50 per replicate) were collected and washed in water to remove food leftovers and yeast. Larvae were placed in drops of ice-cold PBS on the inside of a plastic petri dish lid for dissection. Once the brain was exposed, the brain-lobes were cut out and intact ventral nerve cords (VNC) were collected in a low DNA binding tube containing 100 ul of ice cold Rinaldini’s solution (800mg NaCl, 20mg KCl, 5mg NaH2PO4, 100mg NaHCO3, 100mg glucose in 100ml distilled H2O; filtered through Millipore Steriflip (pluriSelect); 10x can be stored at 4°C). The tubes containing the brains were centrifuged at 300 g for 3min at 4°C. After centrifugation, the supernatant was carefully removed and replaced with freshly made dissociation solution (445ul Scheider’s medium and 50 ul papain (1mg/ml, Sigma-Aldrich, P4762) and 5ul collagenase (1mg/ml, Sigma-Aldrich, C2674)) and incubated for 1hr at 30°C with continuous agitation. Schneider’s culture medium consists of 2.5ml FBS (Thermo Fisher Scientific 16140071), 50μl insulin (Sigma-Aldrich 10516), 500μl pen-strep (Sigma-Aldrich P4458), 2.5ml L-glutamine (Sigma-Aldrich G7513) in 18.925ml Schneider’s medium (Thermo Fisher Scientific 21720024), filtered through Millipore Steriflip (pluriSelect). To guarantee full brain dissociation, the suspension was pipetted up and down every 10 min. After the enzymatic reaction, the samples were centrifuged at 300 g for 3min at 4°C and the supernatant was carefully removed. The dissociated brains were washed with 1ml ice cold Rinaldini’s solution followed by centrifugation at 300 g for 3min at 4°C. After removal of the supernatant, 200ul of Scheider’s medium was added and the tissue was mechanically disrupted by using a 200ul pipette tip for 10-20 times followed by centrifugation at 300 g for 3min at 4°C. The dissociated brains were resuspended in 400ul DPBS with 0.04% BSA followed by filtering through a 10 um pluriStrainer (Cambridge Bioscience, cat. no. 43-50010-03). The filtered suspension was finally centrifuged at 300 g for 3min at 4°C and the tissue was finally resuspended in 43 ul of PBS with 0.04% BSA.

### Single-cell sequencing analysis

Sequencing reads were processed using Cell Ranger (version 6.1.2; 10x Genomics). A custom reference genome was generated using the Cell Ranger mkref pipeline with the *Drosophila melanogaster* reference genome assembly BDGP6.32 and the corresponding gene annotation file Ensembl release 105. The reference FASTA file and GTF annotation file were both obtained from Ensembl. Sequenced libraries were aligned to this reference using Cell Ranger count, generating gene–barcode matrices for downstream analysis.

Raw count matrices were filtered to remove genes with fewer than 3 counts prior to doublet detection. Doublets were identified separately for each sample using scVI-tools SOLO v1.3.0, initialized from a sample-specific scVI model with 1 hidden layer, 20 latent dimensions, and a zero-inflated negative binomial likelihood. Cells with a doublet probability > 0.5 were excluded.

Following doublet removal, singlets were pooled by experimental batch, and adaptive quality-control thresholds were computed independently for each batch using the median absolute deviation (MAD). For the number of detected genes and total UMI counts, lower and upper bounds were set at the median ± 3 MADs of the log1p-transformed distribution, floored at zero. For mitochondrial transcript percentage, an upper bound was set at + 3 MADs of the untransformed distribution. Cells falling outside these batch-specific bounds were excluded. For downstream integration and annotation, batch-corrected latent representations were learned from raw counts using an all-gene scVI model with 1 hidden layer, 128 hidden units, 20 latent dimensions, and a zero-inflated negative binomial likelihood, with experimental batch included as the scVI batch key and replicate and percent mitochondrial content specified as covariates. The resulting scVI latent embedding was used to construct a 15-nearest-neighbour graph and generate the UMAP embeddings. Clustering stability was assessed using Leiden resolutions from 0.2 to 1.2 using pyclustree v0.4.0 and a resolution of 0.4 was selected for downstream analysis. Counts were normalized to 10,000 UMIs per cell and log1p-transformed for marker gene testing with cluster marker genes identified using a one-vs-all Wilcoxon rank-sum test. Cell identities were manually assigned using curated marker gene sets from published studies.

### Differential gene expression analysis

Pseudobulk differential gene expression analysis was performed by summing raw counts across all cells of each annotated cell type within each biological sample to generate one pseudobulk profile per sample and cell type. Differential gene expression analysis was performed in edgeR using a negative binomial generalized linear model with quasi-likelihood inference. Genes with low expression values were filtered using filterByExpr, library sizes were normalized using trimmed mean of M-values normalization, and dispersion was estimated with robust empirical Bayes shrinkage. The design matrix included batch and condition terms, and differential expression between ND-75 KD and Control samples was assessed using a quasi-likelihood F-test on the condition coefficient. P values were adjusted using the Benjamini-Hochberg method, and genes were additionally flagged as potential single-cell artifacts when apparent differential expression was driven by expression in fewer than five cells in the relevant condition. Significant differentially expressed genes were defined using a FDR threshold < 0.05 and an absolute log2 fold change >= 1.0.

### Co-expression gene network analysis

Identification of coordinated gene programs associated with ND-75 Dysfunction beyond individual differentially expressed genes was performed using hdWGCNA v0.4.11 on NSCs. Genes with noisy or non-informative annotations were removed prior to network construction. Genes expressed in at least 5% of cells were used for hdWGCNA, and metacells were generated by grouping cells based on condition and cell-type annotations from the scVI embedding (k = 15, max_shared = 5). A signed NSC co-expression network was constructed with a soft-thresholding power of 9. Module eigengenes and module connectivity (kME) were calculated, and module associations with ND-75 KD were assessed by correlating module eigengenes with condition. Modules with FDR < 0.05 were considered ND-75 KD associated. For integrated prioritization, genes in positively correlated ND-75 KD associated modules were retained if they had a kME >= 0.35 and overlapped with the NSC pseudobulk DEGs.

Functional annotation of hdWGCNA modules was performed using GO enrichment on the top 50 hub genes per module, ranked by kME, with all non-gray module genes used as the background universe. Enrichment was tested with clusterProfiler v4.14.6 and org.Dm.eg.db v3.20.0 across Biological Process (BP), Molecular Function (MF) and Cellular Compartment (CC) ontologies. For Drosophila, as the MF and CC ontologies are sparser compared to BP, ontology-specific thresholds were used: p < 0.05, q < 0.20, minGSSize = 5 for BP and p < 0.10, q < 0.30, minGSSize = 3 for MF and CC. Redundant terms were collapsed using Wang semantic similarity cutoffs of 0.85 for BP and 0.90 for MF and CC. Module labels were assigned from the top non-generic enriched terms and refined manually using GO treeplot hierarchies when needed. NB_M10 was annotated from hub-gene identity because its hub genes were predominantly mtDNA-encoded and GO enrichment returned only indirect annotations.

### Transcription Factor Regulon Analysis

Transcriptional regulators associated with ND-75 KD were identified using pySCENIC v0.12.1 on the NSC subset. Gene regulatory networks were inferred using the FlyBase transcription factor list and pruned against the dm6 motif-ranking database using a motif rank threshold of 3,000 and a normalized enrichment score (NES) threshold of 3.5. Regulon activity was quantified per cell using AUCell with a fixed random seed. Differential regulon activity between control and ND-75 KD NSCs was assessed from regulon AUC scores using two-sided Mann-Whitney U tests, with Benajamini-Hochberg correction for multiple testing. Rank-biserial r was used as the effect size, with positive values indicating higher regulon activity in ND-75 KD NSCs. Identification of regulons associated with the metabolic state was performed using Spearman correlations calculated between regulon AUC scores and Ldh expression overall and separately within each condition. Condition-specific coupling was summarized as *&p* = *p_ND-75_*_KD_ - ^_Control_. The top 40 upregulated regulons ranked by rank-biserial r were visualized as a heatmap with per-cell AUC values z-scored and clipped. Regulons with increased activity and stronger Ldh association in ND-75 KD NSCs were identified by comparing differential regulon activity against Ldh coupling.

### Confocal imaging

Fluorescent images were acquired using a Zeiss LSM880 confocal microscope, with a 63x oil immersion objective, as 12- or 16-bit images. For the larval CNS, we imaged the thoracic segments of the VNC from the ventral side until the neuropil. All images are single sections, unless indicated otherwise. For quantifications, slice thickness was ∼0.8μm, with a 1/3 step size, ensuring complete NSCs were imaged and reconstructed. Images were processed for brightness and contrast using ImageJ.

### Quantifications

For quantification of the mitotic index of NSCs, Dpn-positive NSC on the ventral side of the thoracic VNC at the indicated stage were counted (between 100 and 150). Mitotic index is the number of pH3-positive cells among the Dpn-positive cells. For cell lineage size quantifications, GFP- or tdTomato-labelled clones were generated using the hsFLP/FRT system. Individual lineages derived from a single Dpn-positive NSC were identified by confocal microscopy. Lineage size was quantified by manually counting the total number of GFP- or tdTomato-positive progeny within each clone. Cell counts were performed on confocal z-stack images using Fiji (ImageJ) and the CellCounter plugin. For the EdU incorporation quantifications, the Dpn-positive NSC within each lineage was identified by immunostaining, and the number of EdU-positive progeny was manually counted from confocal z-stack images using Fiji (ImageJ) with the Cell Counter plugin. For brain size, the area of CNS maximum projections was measured. The smFISH quantification was done as described previously ^48^. For quantification of mitochondrial membrane potential, mean fluorescence intensities of tetramethylrhodamine methyl ester (TMRM) were measured using ImageJ software. Regions of interest (ROIs) were drawn around GFP-labelled neural stem cells, and nuclear regions were excluded from the analysis. TMRM intensities were normalized to the mean intensity of control RNAi to determine relative mitochondrial membrane potential. Mean fluorescence intensities of phosphorylated AMPK (pAMPK) were measured using ImageJ software, ROIs were drawn around GFP-labelled NSCs, and pAMPK intensities were normalized to the mean intensity of control RNAi to determine relative pAMPK levels in OxPhos RNAi conditions.

### Deadpanorama Fiji plugin

We developed Deadpanorama, a Fiji ^141^ plugin for easy construction of custom image analysis pipelines for Dpn immunostaining data. Deadpanorama has a modular design allowing segmentation and analysis according to the requirements of an experiment around a robust basic framework. Deadpanorama pipelines begin with segmentation of Dpn labelled nuclei in a user-defined cuboid defined by a selected area and z-slice range. Segmentation uses a frequency bandpass filter followed by agglomerative hierarchical k-means histogram clustering and reconstruction of 3D nuclei based on XY overlap between objects in adjacent z-slices. Trachea fragments area is excluded using a bespoke “intensity flatness” metric defined as the square root of integrated density divided by variance and using a default minimum of 0.2. Deadpanorama integrates CellCounter (https://github.com/fiji/Cell_Counter) by Kurt de Vos to allow manual editing of included nuclei coordinates for counting. Analysis and output modules can be added as required to measure intensity statistics or correlation between chosen marker channels, make positive/negative calls for a specified marker using Kapur’s method ^142^, or give signal ratios between channels in Dpn-positive nuclei or their surrounding area. 3D rendering can be output showing Dpn-positive nuclei marker intensity. The Deadpanorama Eclipse project including pom.xml, source code, compiled classes and .jar files is available on https://github.com/JvdAlab/Drosophila-NSC-mosaicism/tree/main/Deadpanorama.

### Statistical analysis

Statistical analyses were performed in R v4.6.1. One biological replicate is defined as the result of one parental cross. Linear mixed-effects models with brain identity as a random intercept were used for experiments where individual brains contributed multiple clone-level measurements using lme v1.1-37. Ordinary linear models were used for experiments yielding a single measurement per brain. Residual normality and homoscedasticity were assessed using DHARMa v0.5.0, and transforms applied where diagnostics indicated a violation of model assumptions. Variance homogeneity was additionally assessed using the Brown-Forsythe test (car v3.1-3) and a generalized least-squares model with a per-condition variance structure was fitted using nlme v3.1-168 where heteroscedasticity was identified. For factorial designs, Type III tests with sum-to-zero contrasts were applied. Estimated marginal means and pairwise contrasts were computed using emmeans v2.0.4. For small pre-specified contrast sets, familywise error was controlled using multivariate t-distribution adjustment; for large-scale screening comparisons, p-values were adjusted using the Benjamini-Hochberg procedure. Graphs were generated in R and figures assembled in Affinity Designer.

## List of Supplementary Materials

### Supplementary Figures

- Supplementary Figure 1. OxPhos dysfunction rescues proliferation and temporal patterning defects when only a few NSCs are affected.
- Supplementary Figure 2. Gradual increase of the number of NSCs affected with OxPhos dysfunction leads to proliferation impairment.
- Supplementary Figure 3. mtDNA and Complex IV activity are necessary to sustain NSC proliferation
- Supplementary Figure 4. Single-cell RNA-seq quality control, clustering and cell-type annotation
- Supplementary Figure 5. Ldh expression and activity are associated with NSC responses to mitochondrial dysfunction
- Supplementary Figure 6. Transcriptional regulatory networks associated with Ldh expression following Complex I dysfunction
- Supplementary Figure 7. The effects of OxPhos dysfunction extend beyond ATP and ROS production
- Supplementary Figure 8. Glial transporters and gap junctions mediate metabolic support of mitochondrially dysfunctional NSCs
- Supplementary Figure 9. Glial lactate metabolism and lipid metabolic pathways regulate NSC proliferation

### Supplementary Tables

- Supplementary Table 1. Single-cell RNA-seq cell-type marker genes. Provided as separate Excel file.
- Supplementary Table 2. EdgeR pseudo-bulk differential gene expression in each cell-type. Provided as separate Excel file.
- Supplementary Table 3. hdWGCNA module analysis on NSCs Provided as separate Excel file.
- Supplementary Table 4. Gene Ontology analysis of NSC hdGWCNA modules. Provided as separate Excel file.
- Supplementary Table 5. Network analysis of NSC hdGWCNA modules. Provided as separate Excel file.
- Supplementary Table 6. Gene membership of NSC hdGWCNA modules. Provided as separate Excel file.
- Supplementary Table 7. Overlap of differentially expressed genes between edgeR and hdGWCNA in NSCs. Provided as separate Excel file.
- Supplementary Table 8. Transcription factor regulon analysis of differential gene expression in NSCs. Provided as separate Excel file.
- Supplementary Table 9. Drosophila RNAi lines for screening.
- Supplementary Table 10. Larval brain metabolomics. Provided as separate Excel file.
- Supplementary Table 11. Primer sequences used for cloning.

