## Supplementary Figures and Tables S9, S11 for "Tissue-wide metabolic buffering confers resilience to mitochondrial dysfunction"

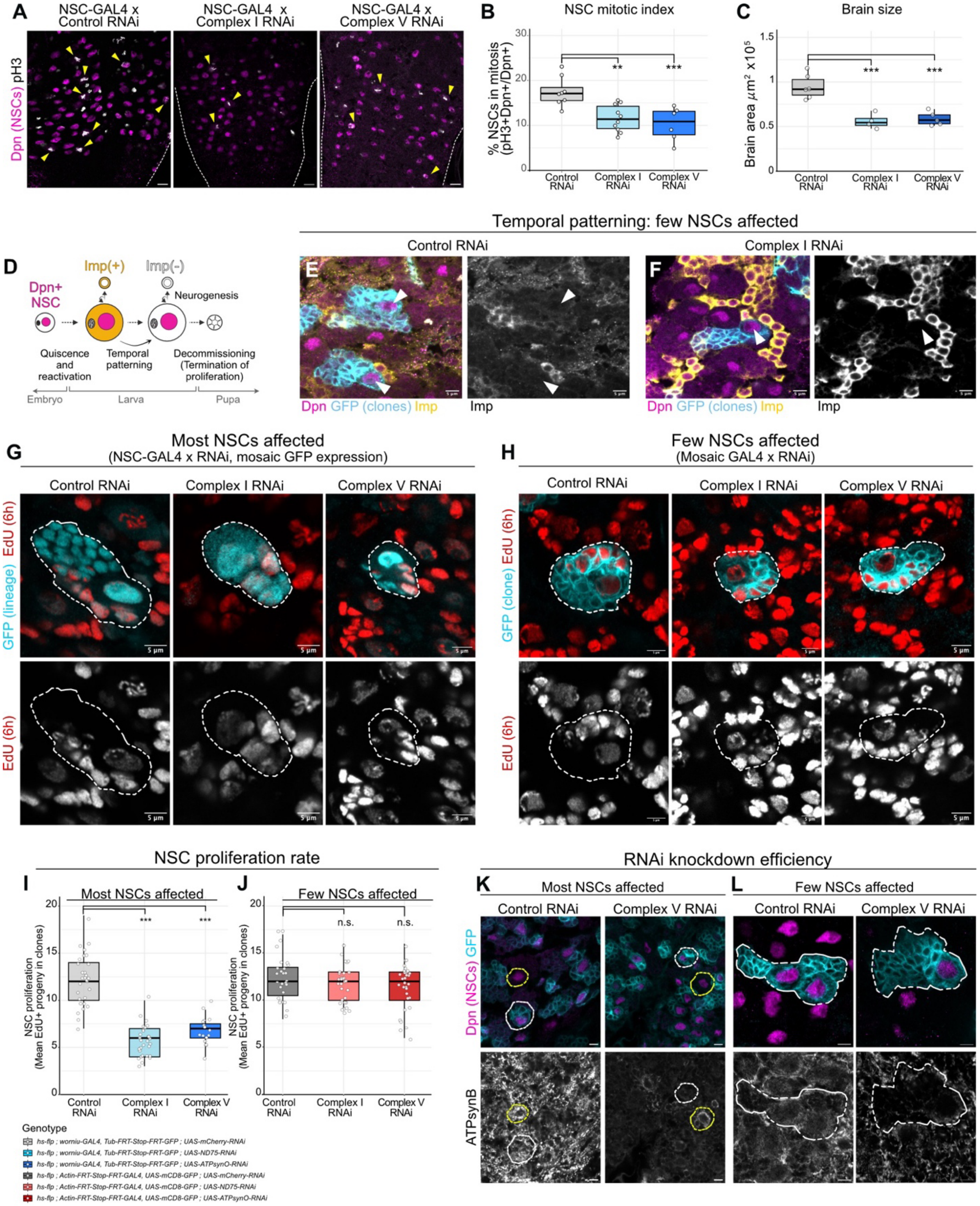


**Supplementary Figure 1. NSC proliferation and temporal patterning upon mosaic OxPhos dysfunction**

(A) phospho-Histone H3 (pH3) and Dpn (NSCs) staining in L3 VNCs expressing control (LexAop-*mCherry*-RNAi), Complex I (LexAOp-*ND75* RNAi), or Complex V (LexAop-*ATPSynO* RNAi) RNAi in NSCs using Dpn-LexA. Dashed lines outline the VNC.
(B,C) Mitotic index of Dpn+ NSCs (B) and brain size (C) from L3 larvae expressing the indicated RNAi constructs in most NSCs (Dpn-LexA(B), worniu-GAL4 (C)). Each datapoint represents an individual brain; Mitotic index: n= 8 for Control RNAi, 10 for Complex I RNAi, 6 for Complex V RNAi; Brain size: n= 6 for Control RNAi, 4 for Complex I RNAi, 5 for Complex V RNAi; unpaired t-tests.
(D) Schematic of the Imp temporal transition in larval NSCs.
(E,F) Imp, Dpn (NSCs) and GFP (clone) staining in L3 VNC clonally expressing control (UAS-*mCherry*-RNAi; E) or Complex I (UAS-*ND75* RNAi; F) RNAi (Actin-FRT-STOP-FRT-GAL4). White arrowheads indicate GFP+ NSCs.
(G,H) EdU and GFP (lineages) staining in VNCs from L3 larvae expressing control, Complex I, or Complex V RNAi in most NSCs with individual NSC lineages randomly labelled by GFP (worniu-GAL4; G), or clonally (Actin-FRT-STOP-FRT-GAL4; H). Dashed lines outline lineages from single NSCs ($Dpn^+$), with EdU+ progeny generated during 6hr EdU feeding.
(I,J) Absolute numbers of newly born cells (EdU+) over a 6h EdU feeding period, per NSC lineage expressing control, Complex I, or Complex V RNAi either specifically in most NSCs (I) or clonally (J). n = 29 clones, 5 brains [Most NSCs affected; Control RNAi], 29 clones, 5 brains [Most NSCs affected; Complex I RNAi], 19 clones, 4 brains [Most NSCs affected; Complex V RNAi], 27 clones, 4 brains [Few NSCs affected; Control RNAi], 31 clones, 5 brains [Few NSCs affected; Complex I RNAi], and 33 clones, 5 brains [Few NSCs affected; Complex V RNAi]; linear mixed effect models.
(K,L) ATPsynB staining in L3 VNC expressing control (*mCherry*-RNAi) or Complex V (*ATPSynO* RNAi) RNAi in most NSCs (worniu-GAL4; K) or clonally (Actin-FRT-STOP-FRT-GAL4; L), with individual NSC lineages marked by GFP. White dashed lines outline NSC lineages expressing the RNAi; yellow dashed lines outline NSCs that are not expressing the RNAi. $p > 0.05$ = n.s.; $p < 0.05$ = *; $p < 0.01$ = **; $p < 0.001$ = *****.** Scale bars: 5 µm (G,H,K,L); 10 µm (A,E,F).

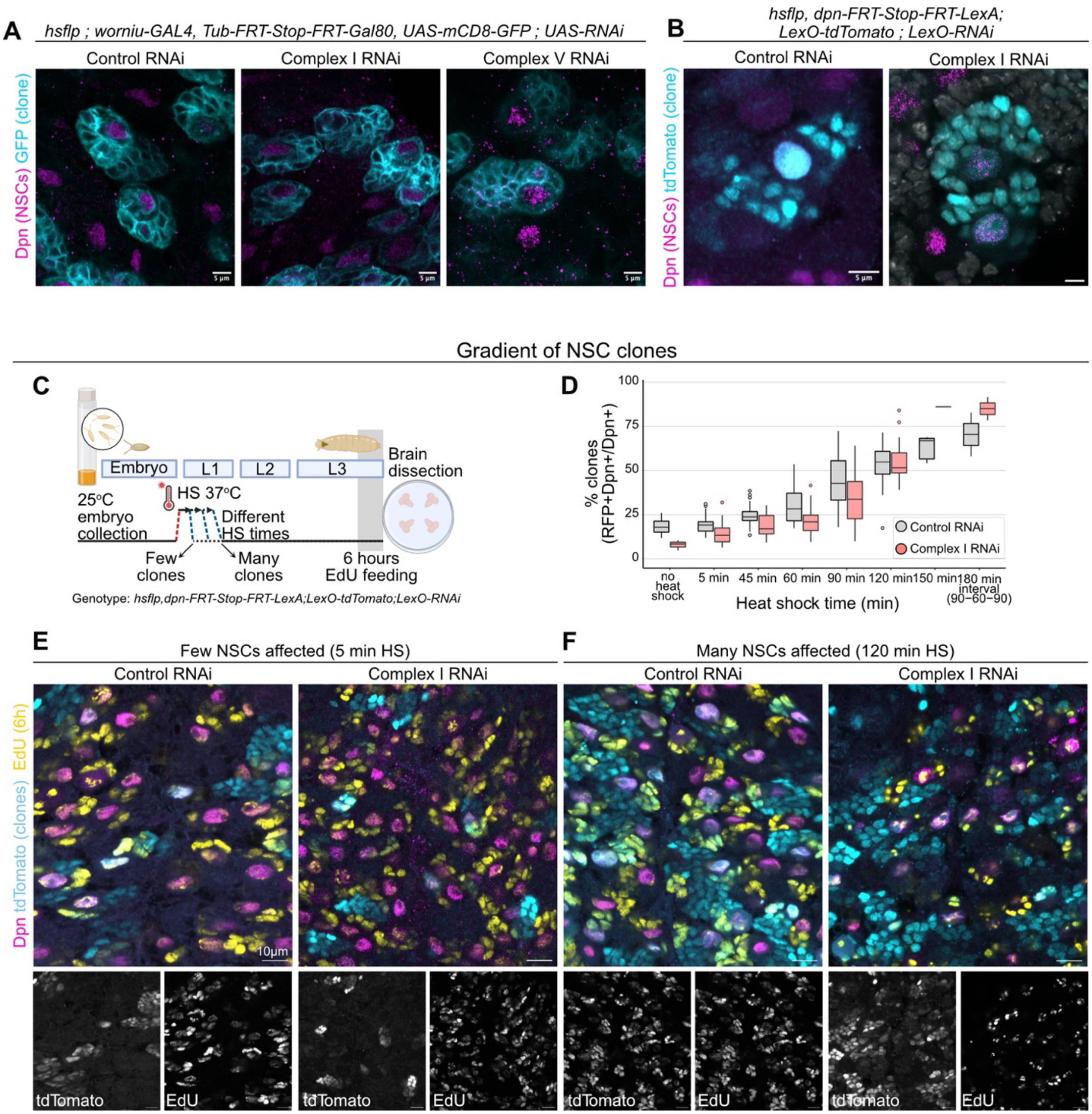


**Supplementary Figure 2. Different types and levels of mosaic OxPhos dysfunction.**
(A,B) Dpn (NSCs) and GFP (A) or tdTomato (B) (clones) staining of L3 VNCs expressing Control, Complex I or Complex V RNAi clonally in NSCs using worniu-GAL4, Tub-FRT-STOP-FRT-GAL80 (A) or Dpn-FRT-STOP-FRT-LexA (B).
(C) Schematic of the heat shock protocol, EdU feeding and brain dissection used for the gradient of clones (related to Figure 1E).
(D) Percentage of Dpn+ NSCs with tdTomato (RFP) expression (clones) per heat shock condition (time in minutes). In the boxplots, the black line shows the median and the coloured boxes the interquartile range between the first and third quartile. 90-60-90: 90 minutes of heat shock, 60 minutes room temperature, 90 minutes heat shock.

(E,F) Dpn (NSCs), tdTomato (clones) and EdU (progeny in last 6 hours) staining in L3 VNCs expressing control (*mCherry*-RNAi) or Complex I (*ND42* RNAi) RNAi clonally in NSCs (Dpn-FRT-STOP-FRT-LexA) after 5 min (E) or 120 min (F) heat shock at L0.
Scale bars: 5 μm (A,B); 10 μm (E,F).

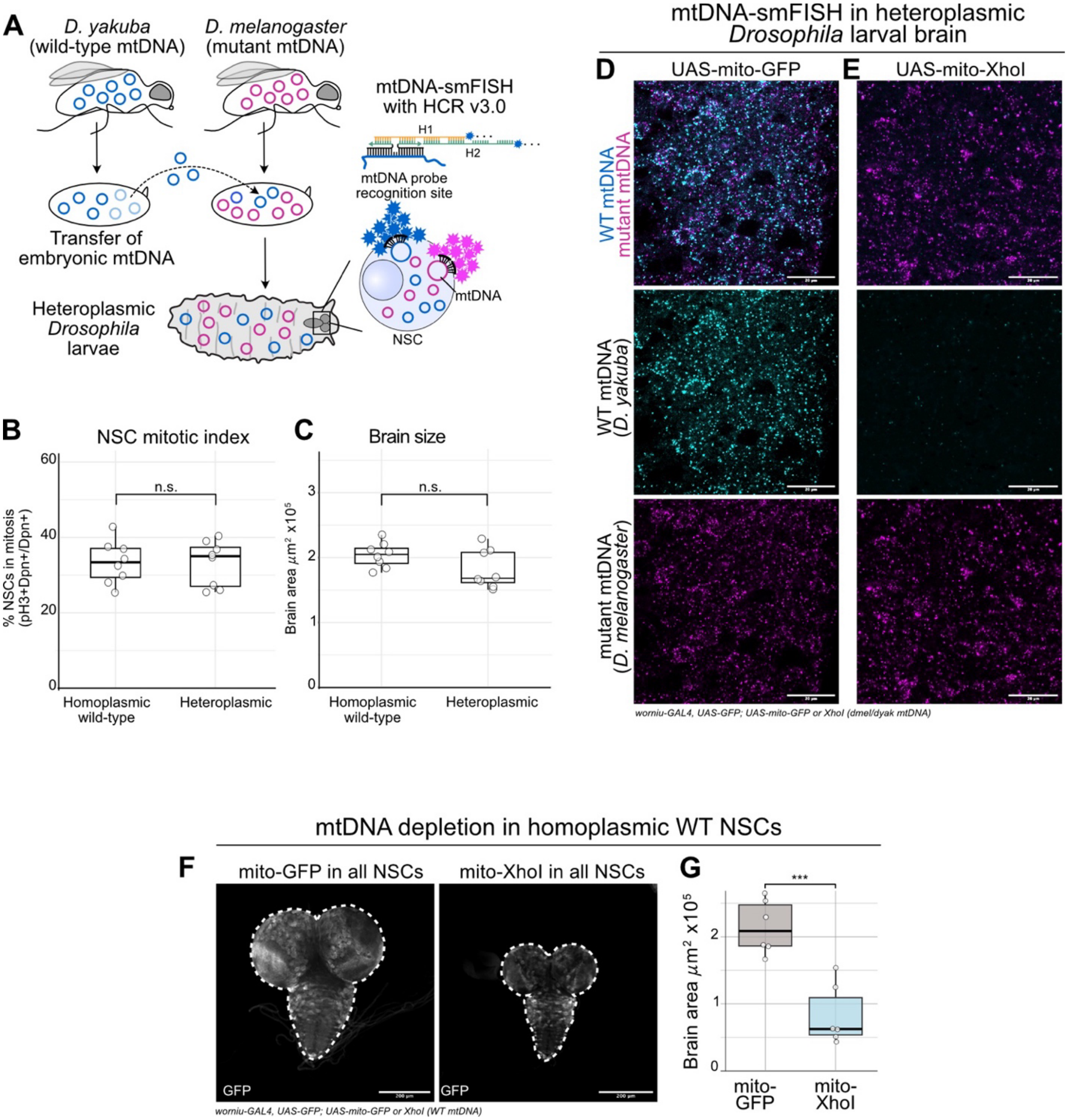


**Supplementary Figure 3. mtDNA and Complex IV activity are required for NSC proliferation**

(A) Generation of heteroplasmic *Drosophila* by interspecies mtDNA transfer between *D. melanogaster* (mutant) and *D. yakuba* (wild type). mtDNA heteroplasmy was detected *in situ* in whole-mount tissues using single-molecule fluorescence *in situ* hybridization of mtDNA (mtDNA-smFISH) with hybridization chain reaction (HCR) for signal amplification, enabling visualization of heteroplasmic mtDNA variants.

(B,C) Mitotic index of $Dpn^{+}$ NSCs (B) and brain size (C) from homoplasmic WT and heteroplasmic L3 larvae. Each datapoint represents an individual brain. n= 8 for homoplasmic WT, 8 for heteroplasmic; unpaired t-tests.

(D,E) mtDNA-smFISH using probes targeting only *D. melanogaster* (mutant, magenta), or only *D.yakuba* (WT, cyan), in heteroplasmic L3 VNCs expressing either mito-GFP (D) or mito-XhoI (E) in most NSCs (worniu-GAL4).
(F,G) Representative images (F) and quantification (G) of brain size of homoplasmic WT L3 brains expressing control (mito-GFP) or mito-XhoI in most NSCs (worniu-GAL4). NSCs lineages are labelled with GFP; dashed lines outline whole brains; data points indicate individual brains; n= 6 for mitoGFP, n= 6 for mitoXhoI; unpaired t-test.
$p > 0.05$ = n.s.; $p < 0.05$ = *; $p < 0.01$ = **; $p < 0.001$ = ***. Scale bars: 20 μm (D,E); 200 μm (F).

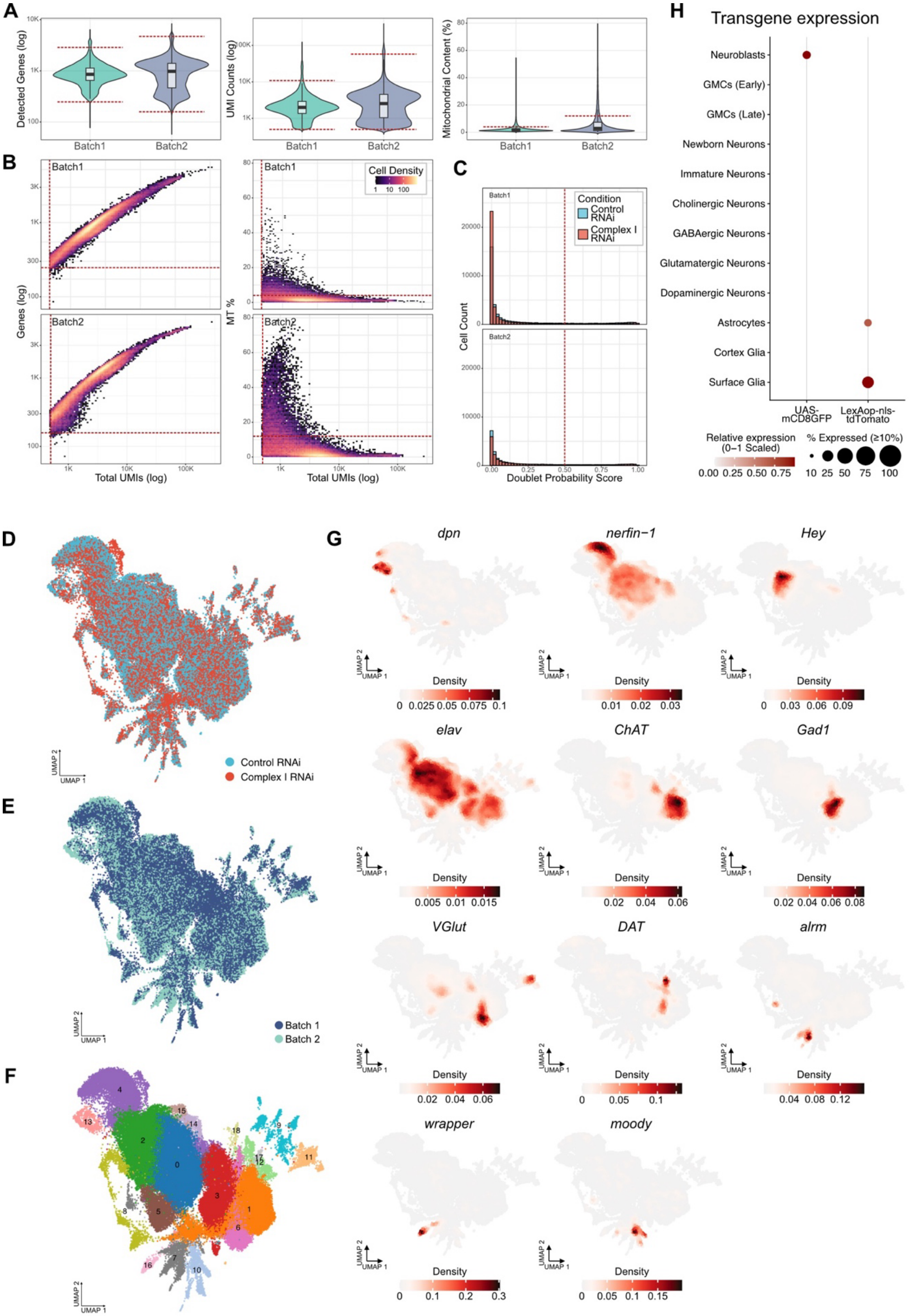
A
B
C
D
E
F
G
H
Detected Genes (log)
UMI Counts (log)
Mitochondrial Content (%)
Batch1
Batch2
Genes (log)
MT %
Total UMIs (log)
Cell Density
Cell Count
Doublet Probability Score
Condition
Control RNAi
Complex I RNAi
Transgene expression
Neuroblasts
GMCs (Early)
GMCs (Late)
Newborn Neurons
Immature Neurons
Cholinergic Neurons
GABAergic Neurons
Glutamatergic Neurons
Dopaminergic Neurons
Astrocytes
Cortex Glia
Surface Glia
UAS-mCD8GFP
LexAop-nls-tdTomato
Relative expression (0–1 Scaled)
% Expressed (≥10%)
Batch 1
Batch 2
UMAP 1
UMAP 2
dpn
nerfin−1
Hey
elav
ChAT
Gad1
VGlut
DAT
alrm
wrapper
moody
Density

**Supplementary Figure 4. Single-cell RNA-seq quality control, clustering and cell-type annotation**

(A) Quality-control distributions for detected genes, total UMI counts, and mitochondrial transcript content across the two scRNA-seq batches. Red dashed lines indicate the lower and upper quality-control thresholds used for filtering cells.

(B) Relationship between total UMI counts and the number of detected genes (left), and between total UMI counts and mitochondrial transcript content (right), shown separately for each batch. Red dashed lines indicate the quality-control thresholds used for cell filtering.

(C) Distribution of SOLO doublet probability scores across samples, for each batch and experimental condition. The red dashed line indicates the quality-control thresholds used for filtering cells.

(D-F) UMAP visualizations of the integrated single-cell transcriptomes, coloured by experimental condition (D), batch (E), or by Leiden cluster identity following clustering at a resolution of 0.4 (F).

(G) Feature-density plots showing the distribution of representative cell-type marker genes across the UMAP embedding, including markers of NSCs (dpn, nerfin-1, Hey), neurons (elav, ChAT, Gad1, VGlut, DAT), and glia (alm, wrapper, moody).

(H) Expression pattern of UAS-mCD8-GFP (worniu-GAL4) and LexO-tdTomato (Repo-LexA) across annotated cell populations. Dot size represents the fraction of expressing cells, and colour intensity indicates relative expression level.

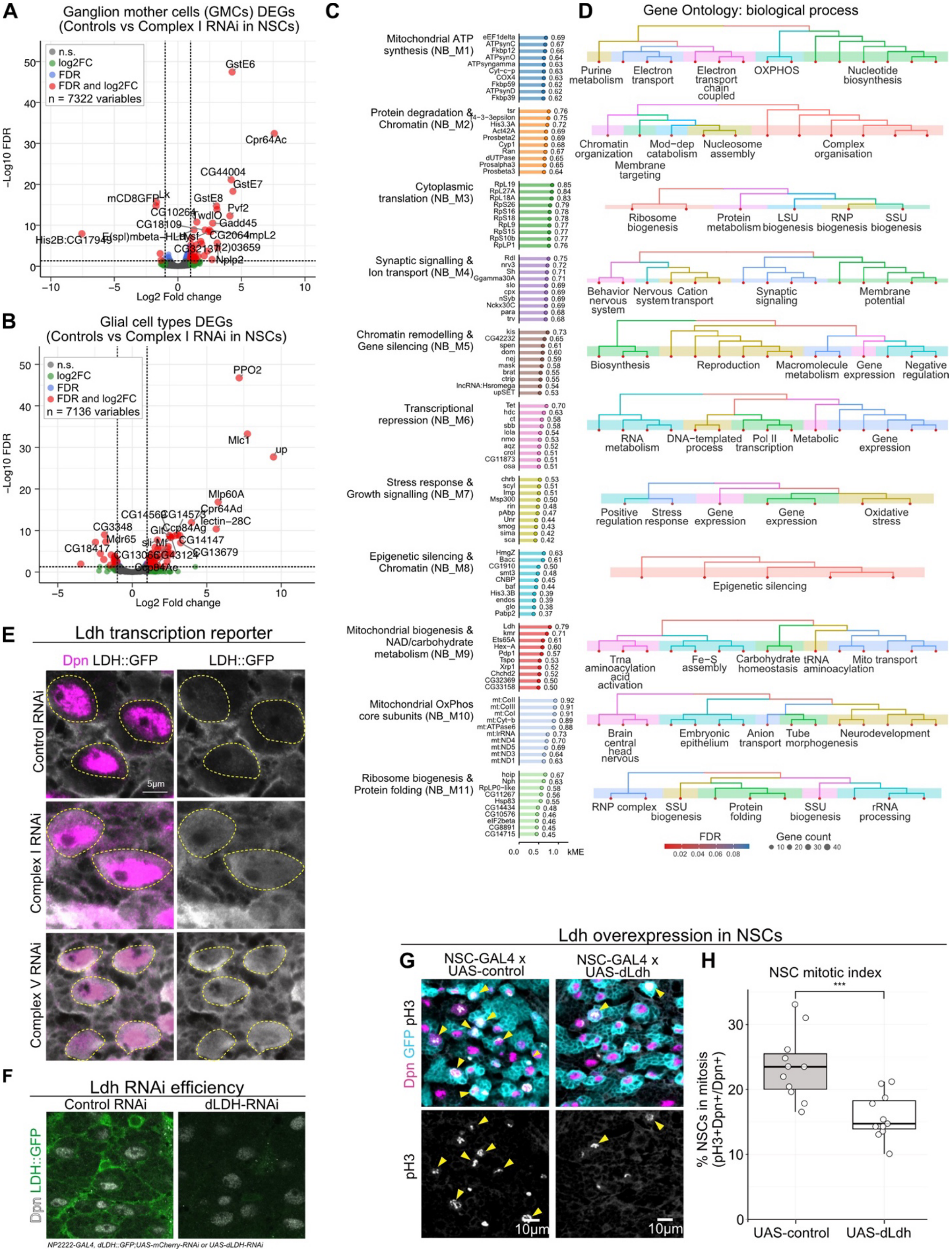
A
Ganglion mother cells (GMCs) DEGs
(Controls vs Complex I RNAi in NSCs)
n.s.
log2FC
FDR
FDR and log2FC
n = 7322 variables
-Log10 FDR
Log2 Fold change
GstE6
Cpr64Ac
CG44004
GstE7
Pvf2
Gadd45
B
Glial cell types DEGs
(Controls vs Complex I RNAi in NSCs)
n.s.
log2FC
FDR
FDR and log2FC
n = 7136 variables
-Log10 FDR
Log2 Fold change
PPO2
Mlc1
up
Mlp60A
Cpr64Ad
C
Mitochondrial ATP synthesis (NB_M1)
Protein degradation & Chromatin (NB_M2)
Cytoplasmic translation (NB_M3)
Synaptic signalling & Ion transport (NB_M4)
Chromatin remodelling & Gene silencing (NB_M5)
Transcriptional repression (NB_M6)
Stress response & Growth signalling (NB_M7)
Epigenetic silencing & Chromatin (NB_M8)
Mitochondrial biogenesis & NAD/carbohydrate metabolism (NB_M9)
Mitochondrial OxPhos core subunits (NB_M10)
Ribosome biogenesis & Protein folding (NB_M11)
kME
D
Gene Ontology: biological process
Purine metabolism
Electron transport
Electron transport chain coupled
OXPHOS
Nucleotide biosynthesis
Chromatin organization
Mod-dep catabolism
Nucleosome assembly
Membrane targeting
Complex organisation
Ribosome biogenesis
Protein metabolism
LSU biogenesis
RNP biogenesis
SSU biogenesis
Behavior nervous system
Nervous system
Cation transport
Synaptic signaling
Membrane potential
Biosynthesis
Reproduction
Macromolecule metabolism
Gene expression
Negative regulation
RNA metabolism
DNA-templated process
Pol II transcription
Metabolic
Gene expression
Positive regulation
Stress response
Gene expression
Gene expression
Oxidative stress
Epigenetic silencing
Trna aminoacylation
Fe-S assembly
Carbohydrate homeostasis
tRNA aminoacylation
Mito transport
acid activation
Brain central head nervous
Embryonic epithelium
Anion transport
Tube morphogenesis
Neurodevelopment
RNP complex
SSU biogenesis
Protein folding
SSU biogenesis
rRNA processing
FDR
Gene count
E
Ldh transcription reporter
Dpn LDH::GFP
LDH::GFP
Control RNAi
Complex I RNAi
Complex V RNAi
5μm
F
Ldh RNAi efficiency
Control RNAi
dLDH-RNAi
Dpn LDH::GFP
NP2222-GAL4, dLDH::GFP;UAS-mCherry-RNAi or UAS-dLDH-RNAi
Ldh overexpression in NSCs
G
NSC-GAL4 x UAS-control
NSC-GAL4 x UAS-dLdh
Dpn GFP pH3
pH3
10μm
10μm
H
NSC mitotic index
***
% NSCs in mitosis (pH3+Dpn+/Dpn+)
UAS-control
UAS-dLdh

**Supplementary Figure 5. Ldh expression and activity are associated with NSC responses to mitochondrial dysfunction**

(A,B) Pseudobulk differential gene expression (edgeR) between control and mitochondrial dysfunction (Complex I RNAi) conditions in (A) Ganglion mother Cells (GMCs) and (B) combined glial cell types. Significant differentially expressed genes defined as FDR $< 0.05$ and absolute log2 fold change $\geq 1$. Colours indicate genes passing the FDR threshold, log2 fold-change threshold, both thresholds, or neither.

(C) Hub genes from NSC gene co-expression modules (hdWGCNA). Genes are grouped by module and ranked according to module connectivity (kME), shown on the x-axis. Module labels indicate representative biological processes associated with each gene co-expression module.

(D) Gene Ontology Biological Process enrichment of hdWGCNA modules. Tree plots show relationships among enriched GO terms, with related terms grouped by semantic similarity. Node colour represents FDR and node size represents gene count.

(E) *Ldh* transcriptional reporter expression and Dpn immunostaining in L3 VNCs expressing control, Complex I, or Complex V RNAi in most NSCs (worniu-GAL4). The yellow outlines delineating NSC regions.

(F) Dpn and GFP immunostaining in L3 VNCs expressing control (mCherry RNAi) or dLDH RNAi in combination with an LDH::GFP reporter line in cortex glia (NP2222-GAL4).

(G) Phospho-Histone H3 (pH3) and Dpn immunostaining in L3 VNCs overexpressing either UAS-GFP or UAS-dLDH in most NSCs (worniu-GAL4). NSCs lineages are marked with GFP. The yellow arrows indicate pH3+ NSCs.

(H) Mitotic index of $Dpn^{+}$ NSCs from L3 larvae expressing the indicated constructs in NSCs.
Data points indicate individual brains, n= 11 for Control and 11 for dLdh; unpaired t-test.
$p > 0.05$ = n.s.; $p < 0.05$ = *; $p < 0.01$ = **; $p < 0.001$ = ***. Scale bars: 5 µm (E,F), 10 µm (G).

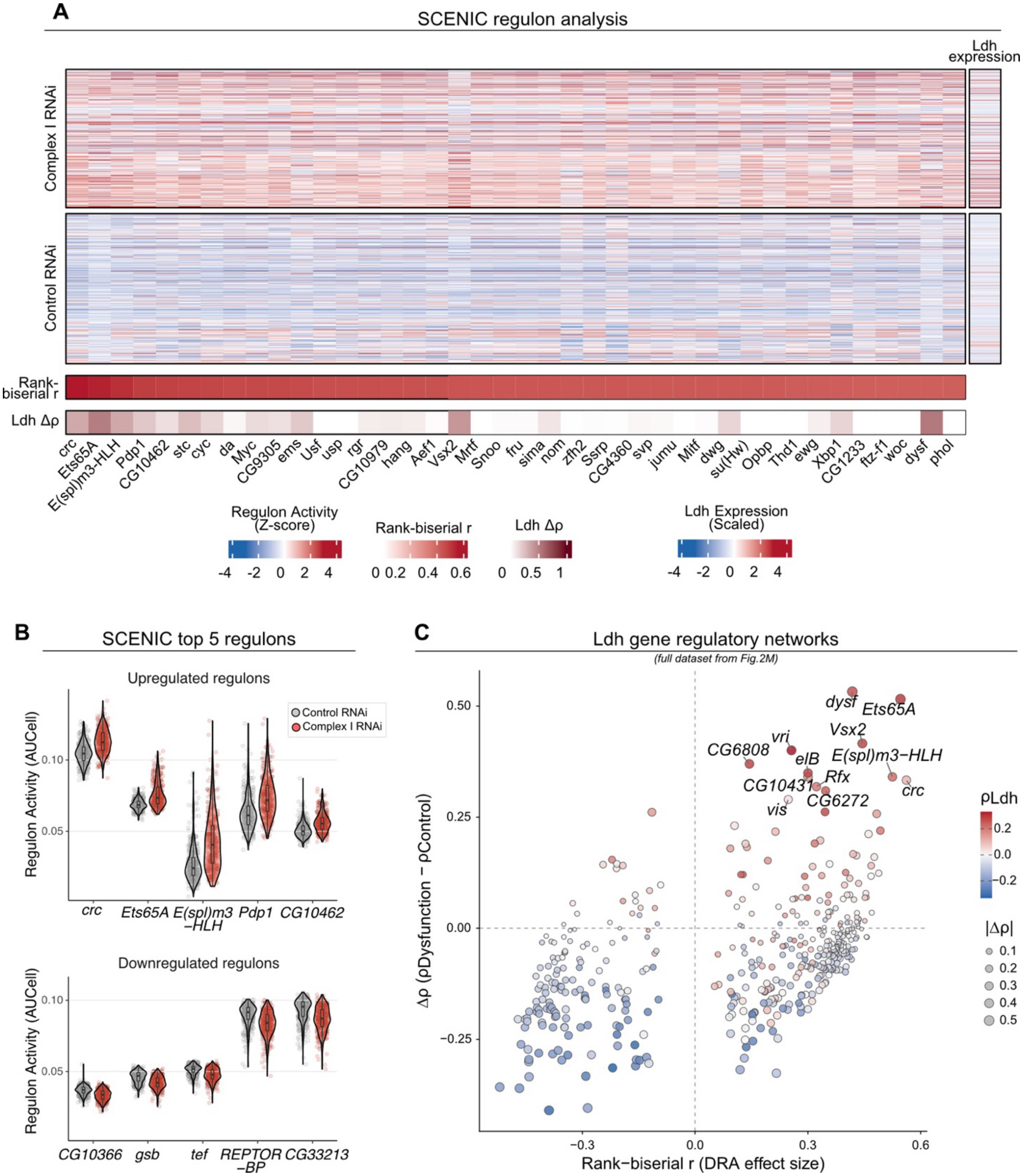


**Supplementary Figure 6. Transcriptional regulatory networks associated with Complex I dysfunction**

(A) Per-cell activity scores of top 40 upregulated transcription factor regulons in NSCs identify by pySCENIC, ranked by differential regulon activity between Complex I RNAi and Control. Regulon activity is shown as per-regulon z-scored AUCell values, with cells grouped by condition. Z-scored *Ldh* expression across NSCs shown alongside each cell. Bottom annotations indicate the rank-biserial effect size for differential regulon activity and the change in *Ldh* coupling for each regulon. For visualization, negative $\Delta\rho$ values are displayed as zero in the bottom annotation.

(B) Relationship between differential regulon activity and *Ldh* coupling across NSC regulons. The x-axis shows the rank-biserial effect size for differential regulon activity between Complex I RNAi and Control; y-axis shows the change in Spearman correlation between regulon activity and *Ldh* expression. Colour indicates overall Spearman correlation between regulon activity and *Ldh* expression. For significantly differentially active regulons (FDR $< 0.05$), point size is proportional to $|\Delta\rho|$; non-significant regulons are shown at a fixed size and lower opacity. Selected significantly upregulated regulons with the largest increases in *Ldh* coupling are labelled

(C) Per-cell activity of the five most up- and down-regulated transcription factor regulons in NSCs, ranked by differential regulon activity between Complex I and Control RNAi. Regulon activity quantified by pySCENIC AUCell score. Points represent individual cells; violin width indicates the distribution of regulon activity; embedded boxplots show the median and interquartile range. Control RNAi in grey, Complex I RNAi in red.

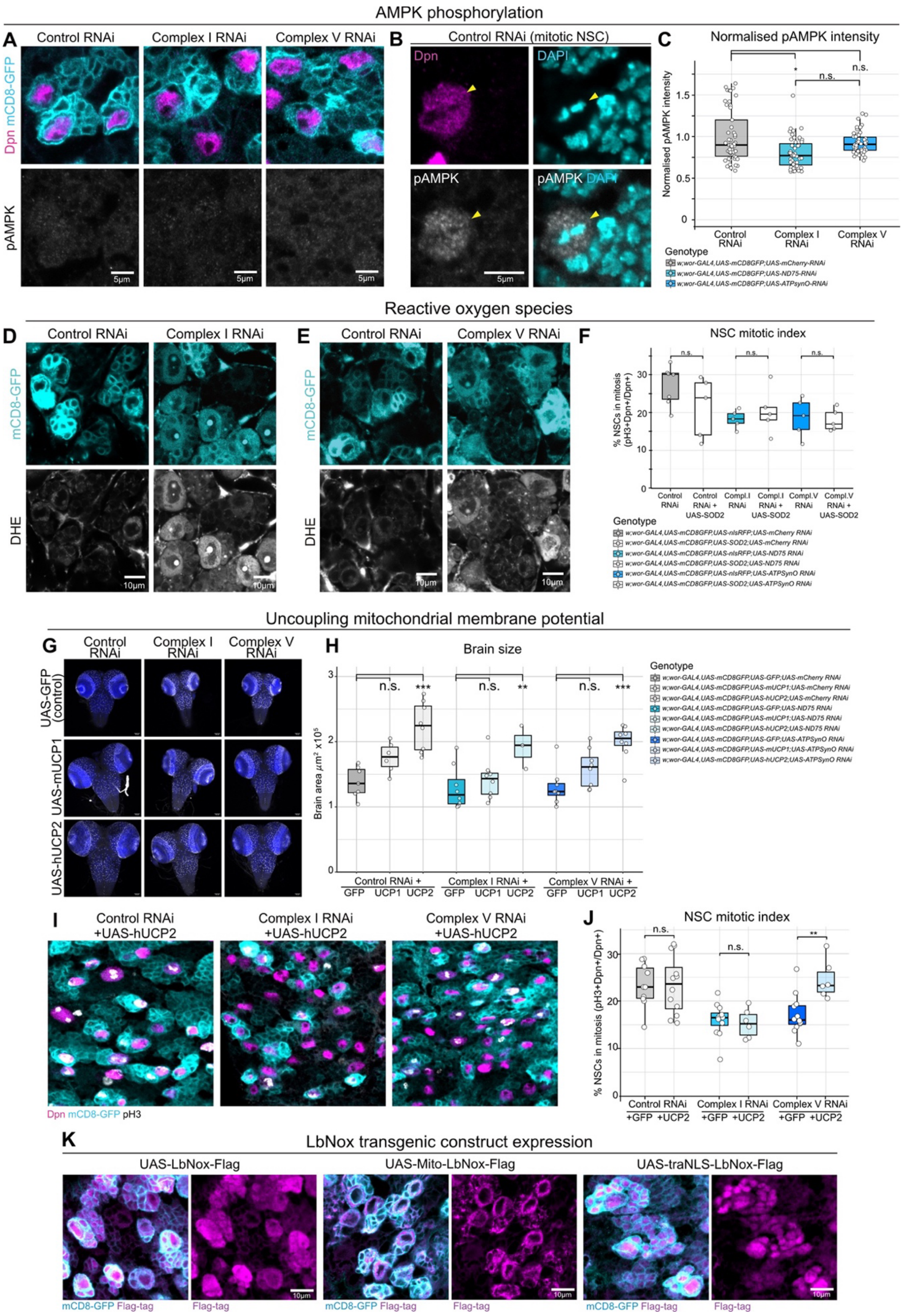

AMPK phosphorylation
A
Control RNAi
Complex I RNAi
Complex V RNAi
Dpn mCD8-GFP
pAMPK
5µm
B
Control RNAi (mitotic NSC)
Dpn
DAPI
pAMPK
pAMPK DAPI
C
Normalised pAMPK intensity
Normalised pAMPK intensity
n.s.
n.s.
Control RNAi
Complex I RNAi
Complex V RNAi
Genotype
w;wor-GAL4,UAS-mCD8GFP;UAS-mCherry-RNAi
w;wor-GAL4,UAS-mCD8GFP;UAS-ND75-RNAi
w;wor-GAL4,UAS-mCD8GFP;UAS-ATPsynO-RNAi
Reactive oxygen species
D
Control RNAi
Complex I RNAi
mCD8-GFP
DHE
10µm
E
Control RNAi
Complex V RNAi
F
NSC mitotic index
% NSCs in mitosis (pH3+Dpn+/Dpn+)
n.s.
Control RNAi
Control RNAi + UAS-SOD2
Compl.I RNAi
Compl.I RNAi + UAS-SOD2
Compl.V RNAi
Compl.V RNAi + UAS-SOD2
Genotype
w;wor-GAL4,UAS-mCD8GFP,UAS-nlsRFP;UAS-mCherry RNAi
w;wor-GAL4,UAS-mCD8GFP,UAS-SOD2;UAS-mCherry RNAi
w;wor-GAL4,UAS-mCD8GFP,UAS-nlsRFP;UAS-ND75 RNAi
w;wor-GAL4,UAS-mCD8GFP,UAS-SOD2;UAS-ND75 RNAi
w;wor-GAL4,UAS-mCD8GFP,UAS-nlsRFP;UAS-ATPSynO RNAi
w;wor-GAL4,UAS-mCD8GFP,UAS-SOD2;UAS-ATPSynO RNAi
Uncoupling mitochondrial membrane potential
G
Control RNAi
Complex I RNAi
Complex V RNAi
UAS-GFP (control)
UAS-mUCP1
UAS-hUCP2
H
Brain size
Brain area µm² x10⁵
n.s.
***
**
Control RNAi +
Complex I RNAi +
Complex V RNAi +
GFP UCP1 UCP2
Genotype
w;wor-GAL4,UAS-mCD8GFP,UAS-GFP;UAS-mCherry RNAi
w;wor-GAL4,UAS-mCD8GFP,UAS-mUCP1;UAS-mCherry RNAi
w;wor-GAL4,UAS-mCD8GFP,UAS-hUCP2;UAS-mCherry RNAi
w;wor-GAL4,UAS-mCD8GFP,UAS-GFP;UAS-ND75 RNAi
w;wor-GAL4,UAS-mCD8GFP,UAS-mUCP1;UAS-ND75 RNAi
w;wor-GAL4,UAS-mCD8GFP,UAS-hUCP2;UAS-ND75 RNAi
w;wor-GAL4,UAS-mCD8GFP,UAS-GFP;UAS-ATPSynO RNAi
w;wor-GAL4,UAS-mCD8GFP,UAS-mUCP1;UAS-ATPSynO RNAi
w;wor-GAL4,UAS-mCD8GFP,UAS-hUCP2;UAS-ATPSynO RNAi
I
Control RNAi +UAS-hUCP2
Complex I RNAi +UAS-hUCP2
Complex V RNAi +UAS-hUCP2
Dpn mCD8-GFP pH3
J
NSC mitotic index
% NSCs in mitosis (pH3+Dpn+/Dpn+)
n.s.
**
Control RNAi
Complex I RNAi
Complex V RNAi
+GFP +UCP2
K
LbNox transgenic construct expression
UAS-LbNox-Flag
UAS-Mito-LbNox-Flag
UAS-traNLS-LbNox-Flag
mCD8-GFP Flag-tag
Flag-tag
10µm

**Supplementary Figure 7. The effects of OxPhos dysfunction extend beyond ATP and ROS production**

(A,B) Phosphorylated AMP-activated protein kinase (pAMPK), Dpn (NSCs) and GFP (NSC lineages) staining in L3 VNCs expressing control (UAS-*mCherry*-RNAi), Complex I (UAS-*ND75* RNAi) or Complex V (UAS-ATPsynO RNAi) RNAi in most NSCs (worniu-GAL4). A mitotic NSC (B) expressing control RNAi (UAS-*mCherry*-RNAi) in NSCs as positive control for pAMPK staining, with DAPI as nuclear stain.

(C) Quantification of normalised pAMPK intensity expressing the indicated constructs in NSCs. Each datapoint represents an individual brain, n = 12 [Control RNAi], 14 [Complex I RNAi], 13 [Complex V RNAi]; linear mixed effect models.

(D,E) Dihydroethidium (DHE) staining in L3 VNCs expressing control (UAS-*mCherry*-RNAi; D,E), Complex I RNAi (UAS-*ND75* RNAi; D) or Complex V RNAi (UAS-*ATPsynO* RNAi; E), in most NSCs (worniu-GAL4). The NSCs lineages are labelled with GFP (cyan).

(F) Mitotic index of $Dpn^+$ NSCs from L3 larvae expressing Control (UAS-mCherry RNAi), Complex I (UAS-ND75 RNAi) or Complex V (UAS-ATPsynO RNAi) RNAi with or without UAS-SOD2 overexpression in NSCs (worniu-Gal4). Each datapoint represents an individual brain, n = 7 [Control RNAi], 5 [Control RNAi; SOD2], 5 [Complex I RNAi], 5 [Complex I RNAi; SOD2], 5 [Complex V RNAi], 5 [Complex V RNAi; SOD2]; linear mixed effect models.

(G) L3 brains expressing control, Complex I or Complex V RNAi in most NSCs while overexpressing either mUCP1 or hUCP2.

(H) Quantification of L3 brain size expressing the indicated genotypes. Data points indicate individual brains, n=7 [Control RNAi, GFP], 6 [Control RNAi; UCP1], 8 [Control RNAi; UCP2], 8 [Complex I RNAi, GFP], 8 [Complex I RNAi, UCP1], 3 [Complex I RNAi, UCP2], 8 [Complex V RNAi; GFP], 8 [Complex V RNAi; UCP1], 8 [Complex V RNAi; UCP2]; Dunnett's test.

(I,J) Phospho-Histone H3 (pH3), Dpn (NSCs) and GFP (NSC lineages) immunostaining (I) and mitotic index (J) in L3 VNCs expressing Control (UAS-mCherry RNAi), Complex I (UAS-ND75 RNAi) or Complex V (UAS-ATPsynO RNAi) RNAi in most NSCs (worniu-GAL4), while overexpressing UAS-hUCP2. Each datapoint represents an individual brain, n = 10 [Control RNAi; GFP], 12 [Control RNAi, UCP2], 12 [Complex I RNAi; GFP], 6 [Complex I RNAi; UCP2], 12 [Complex V RNAi; GFP], 6 [Complex V RNAi; UCP2]; linear mixed effect models.

(K) Dpn (NSCs), GFP (NSC lineages) and FLAG (LbNox transgene) immunostaining in L3 VNCs expressing UAS-LbNox, UAS-traNLS-LbNox or mito-LbNox in most NSCs (worniu-GAL4).

$p > 0.05$ = n.s.; $p < 0.05$ = *; $p < 0.01$ = **; $p < 0.001$ = ***. Scale bars: 5 μm (A,B), 10 μm (D,E,I,K).

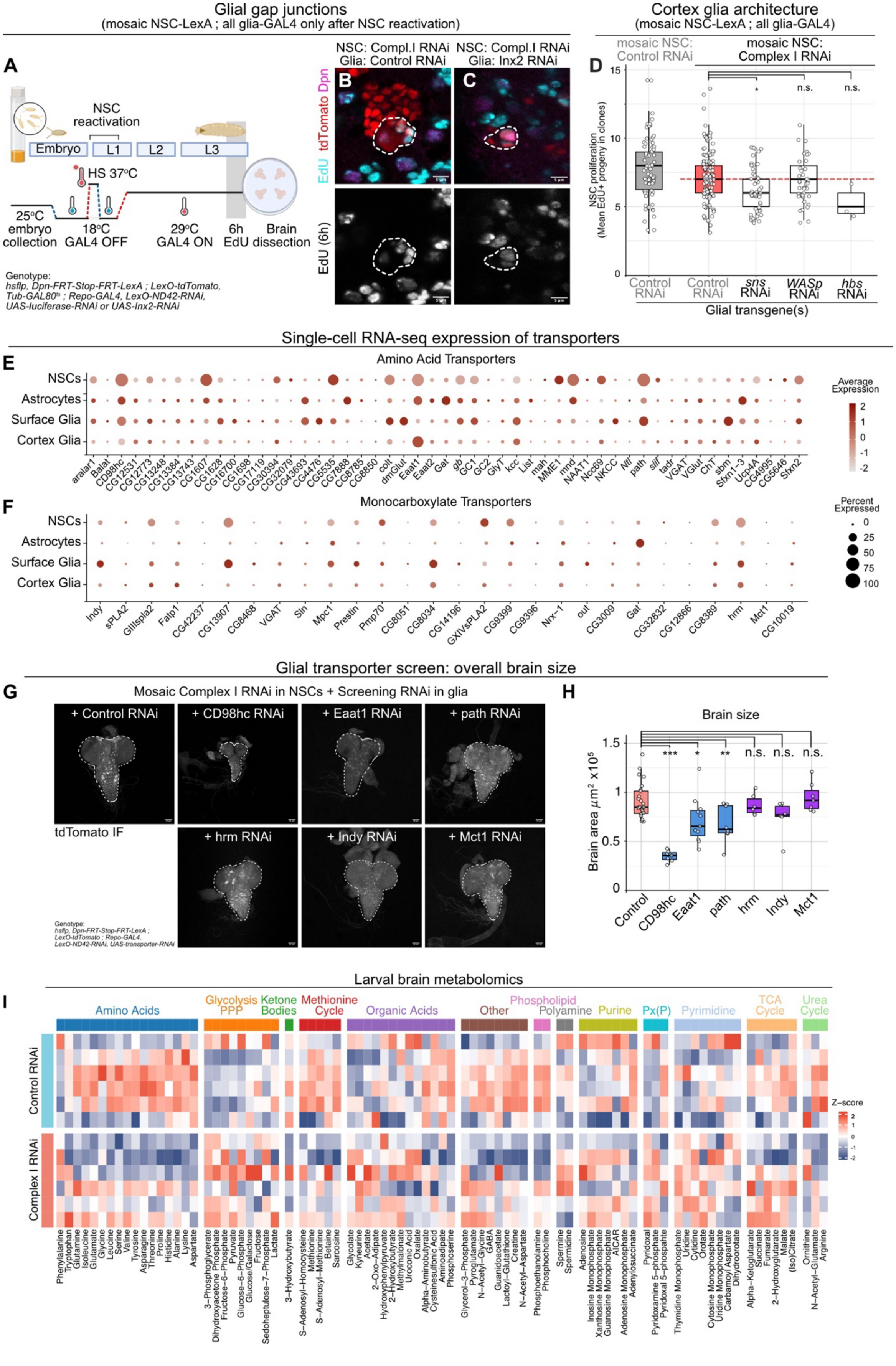
Glial gap junctions
(mosaic NSC-LexA ; all glia-GAL4 only after NSC reactivation)
A
NSC reactivation
Embryo
L1
L2
L3
HS 37°C
25°C embryo collection
18°C GAL4 OFF
29°C GAL4 ON
6h EdU
Brain dissection
Genotype:
hsflp, Dpn-FRT-Stop-FRT-LexA ; LexO-tdTomato, Tub-GAL80ts ; Repo-GAL4, LexO-ND42-RNAi, UAS-luciferase-RNAi or UAS-Inx2-RNAi
NSC: Compl.I RNAi Glia: Control RNAi
NSC: Compl.I RNAi Glia: Inx2 RNAi
B
C
EdU tdTomato Dpn
EdU (6h)
Cortex glia architecture
(mosaic NSC-LexA ; all glia-GAL4)
mosaic NSC: Control RNAi
mosaic NSC: Complex I RNAi
D
NSC proliferation (Mean EdU+ progeny in clones)
*
n.s.
n.s.
Control RNAi
Control RNAi
sns RNAi
WASp RNAi
hbs RNAi
Glial transgene(s)
Single-cell RNA-seq expression of transporters
E
Amino Acid Transporters
NSCs
Astrocytes
Surface Glia
Cortex Glia
aralar1
Balat
CD98hc
CG12531
CG12773
CG13248
CG13384
CG13743
CG1607
CG1628
CG16700
CG1698
CG17119
CG30394
CG32079
CG43693
CG4476
CG5535
CG7888
CG8785
CG8850
colt
dmGlut
Eaat1
Eaat2
Gat
gb
GC1
GC2
GlyT
kcc
List
mah
MME1
mnd
NAAT1
Nckx30
NKCC
Ntl
path
slif
tadr
VGAT
VGlut
ChT
sbm
Sfxn1-3
Ucp4A
CG4995
CG5646
Sfxn2
Average Expression
F
Monocarboxylate Transporters
Indy
sPLA2
GIIIspla2
Fatp1
CG42237
CG13907
CG8468
VGAT
Sln
Mpc1
Prestin
Pmp70
CG8051
CG8034
CG14196
GXIVsPLA2
CG9399
CG9396
Nrx-1
out
CG3009
Gat
CG32832
CG12866
CG8389
hrm
Mct1
CG10019
Percent Expressed
Glial transporter screen: overall brain size
G
Mosaic Complex I RNAi in NSCs + Screening RNAi in glia
+ Control RNAi
+ CD98hc RNAi
+ Eaat1 RNAi
+ path RNAi
tdTomato IF
+ hrm RNAi
+ Indy RNAi
+ Mct1 RNAi
Genotype:
hsflp, Dpn-FRT-Stop-FRT-LexA ; LexO-tdTomato ; Repo-GAL4, LexO-ND42-RNAi, UAS-transporter-RNAi
H
Brain size
Brain area μm² x10⁵
***
*
**
n.s.
n.s.
n.s.
Control
CD98hc
Eaat1
path
hrm
Indy
Mct1
I
Larval brain metabolomics
Amino Acids
Glycolysis PPP
Ketone Bodies
Methionine Cycle
Organic Acids
Other
Phospholipid
Polyamine
Purine
Px(P)
Pyrimidine
TCA Cycle
Urea Cycle
Control RNAi
Complex I RNAi
Z-score
Phenylalanine
Tryptophan
Glutamine
Isoleucine
Glutamate
Glycine
Leucine
Serine
Valine
Tyrosine
Asparagine
Threonine
Proline
Histidine
Alanine
Lysine
Aspartate
3-Phosphoglycerate
Dihydroxyacetone Phosphate
Fructose-6-Phosphate
Pyruvate
Glucose-6-Phosphate
Glucose/Galactose
Fructose
Sedoheptulose-7-Phosphate
Lactate
3-Hydroxybutyrate
S-Adenosyl-Homocysteine
Methionine
S-Adenosyl-Methionine
Betaine
Sarcosine
Glycolate
Kyneurine
Acetate
2-Oxo-Adipate
Hydroxyphenylpyruvate
2-Hydroxybutyrate
Methylmalonate
Urocanic Acid
Oxalate
Alpha-Aminobutyrate
Cysteinesulfonic Acid
Aminoadipate
Phosphoserine
Glycerol-3-Phosphate
Pyroglutamate
N-Acetyl-Glycine
GABA
Guanidoacetate
Lactoyl-Glutathione
Creatine
N-Acetyl-Aspartate
Phosphoethanolamine
Phosphocholine
Spermine
Spermidine
Adenosine
Inosine Monophosphate
Xanthosine Monophosphate
Guanosine Monophosphate
AICAR
Adenosine Monophosphate
Adenylosuccinate
Pyridoxal
Pyridoxamine 5-phosphate
Pyridoxal 5-phosphate
Thymidine Monophosphate
Uridine
Cytidine
Orotate
Cytosine Monophosphate
Uridine Monophosphate
Carbamoyl Aspartate
Dihydroorotate
Alpha-Ketoglutarate
Succinate
Fumarate
2-Hydroxyglutarate
Malate
(Iso)Citrate
Ornithine
N-Acetyl-Glutamate
Arginine

**Supplementary Figure 8. Glial transporters and gap junctions are required to support proliferation of NSCs with OxPhos dysfunction**

(A) Schematic of the heat shock, temperature shifting and EdU feeding protocol for the glial gap junction RNAi experiment (see Figure 4C).

(B,C) EdU (NSC progeny, 6hr feeding), Dpn (NSCs) and tdTomato (NSC lineages) immunostaining in L3 VNCs with clonal Complex I RNAi expression in NSCs (Dpn-FRT-STOP-FRT-LexA), while expressing either control (UAS-luciferase RNAi, B) or UAS-inx2 RNAi in the surrounding glial cells (Repo-GAL4). Dashed lines outline lineages from single NSCs ($Dpn^{+}$).

(D) NSC proliferation after knocking down genes involved in cortex glia architecture in the glia (Repo-GAL4), with clonal expression of Complex I RNAi in NSCs. Absolute numbers of newly produced cells (EdU+) per NSC lineage in the VNC with the indicated genotypes. Each datapoint represents number of clones within individual brains, n = 1413 clones, 321 brains in total; linear mixed effect models.

(E,F) Single-cell RNA-seq expression of amino acid (E) and monocarboxylate (F) transporter genes in NSCs, astrocytes, surface glia and cortex glia. Dot size represents the percentage of cells expressing each gene; colour intensity represents scaled average expression within each cell type.

(G,H) L3 brains (G) and brain size quantification (H) expressing control, CD98hc, Eaat1, path, hrm, Indy or Mct1 RNAi in the glia (Repo-GAL4), while NSCs are clonally expressing Complex I RNAi (LexAop-ND42 RNAi). The dashed lines outline the whole brains. Each datapoint represents number of clones within individual brains, n = 25 [Control RNAi], 7 [CD98hc RNAi], 11 [Eaat1 RNAi], 8 [path RNAi], 10 [hrm RNAi], 6 [Indy RNAi], 7 [Mct1 RNAi]; Dunnett's test.

(I) Relative metabolite abundance in whole L3 brains expressing Control or Complex I RNAi in most NSCs (worniu-GAL4). Columns represent individual metabolites grouped by metabolic pathway, and rows represent independent biological replicates. Metabolite abundances are shown as Z-scores calculated across samples for each metabolite, with blue and red indicating lower and higher relative abundance, respectively. Pathway categories are indicated by coloured bars above the heatmap.

$p > 0.05$ = n.s.; $p < 0.05$ = *; $p < 0.01$ = **; $p < 0.001$ = ***. Scale bars are 5 μm (B,C), 50 μm (G).

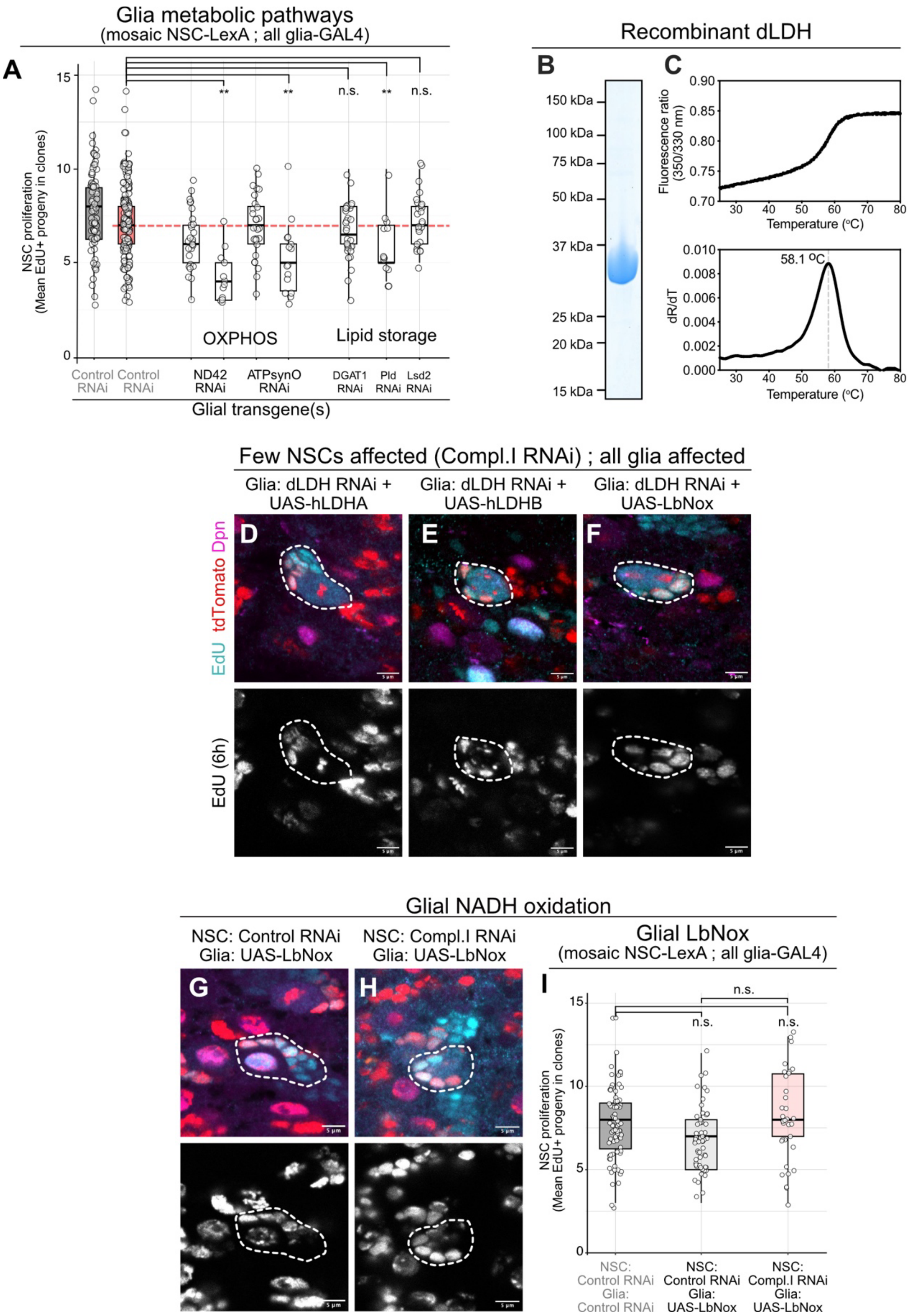
Glia metabolic pathways
(mosaic NSC-LexA ; all glia-GAL4)
A
NSC proliferation
(Mean EdU+ progeny in clones)
15
10
5
0
** ** n.s. ** n.s.
OXPHOS
Lipid storage
Control RNAi
Control RNAi
ND42 RNAi
ATPsynO RNAi
DGAT1 RNAi
Pld RNAi
Lsd2 RNAi
Glial transgene(s)
Recombinant dLDH
B
150 kDa
100 kDa
75 kDa
50 kDa
37 kDa
25 kDa
20 kDa
15 kDa
C
Fluorescence ratio (350/330 nm)
0.90
0.85
0.80
0.75
0.70
30 40 50 60 70 80
Temperature (°C)
58.1 °C
dR/dT
0.010
0.008
0.006
0.004
0.002
0.000
30 40 50 60 70 80
Temperature (°C)
Few NSCs affected (Compl.I RNAi) ; all glia affected
Glia: dLDH RNAi + UAS-hLDHA
Glia: dLDH RNAi + UAS-hLDHB
Glia: dLDH RNAi + UAS-LbNox
D
E
F
EdU tdTomato Dpn
EdU (6h)
Glial NADH oxidation
NSC: Control RNAi
Glia: UAS-LbNox
NSC: Compl.I RNAi
Glia: UAS-LbNox
G
H
Glial LbNox
(mosaic NSC-LexA ; all glia-GAL4)
I
n.s.
n.s.
n.s.
NSC proliferation
(Mean EdU+ progeny in clones)
15
10
5
0
NSC: Control RNAi Glia: Control RNAi
NSC: Control RNAi Glia: UAS-LbNox
NSC: Compl.I RNAi Glia: UAS-LbNox

**Supplementary Figure 9. Glial lactate metabolism and lipid metabolic pathways regulate NSC proliferation**

(A) NSC proliferation after knocking down genes involved in OxPhos or Lipid storage in the glia (Repo-GAL4), where NSCs are clonally expressing Complex I RNAi. Absolute numbers of newly synthesised cells (EdU+, 6hr feeding) per NSC lineage in the VNC with the indicated genotypes. Each datapoint represents number of clones within individual brains, n = 1413 clones, 321 brains in total; linear mixed effect models.

(B,C) PAGE (B) and nanoDSF (C), to validate *Drosophila* LDH (dLDH) recombinant protein expression.

(D-F) EdU (NSC progeny, 6hr feeding), Dpn (NSCs) and tdTomato (NSC lineages) immunostaining in L3 VNCs with clonal Complex I RNAi in NSCs (Dpn-FRT-STOP-FRT-LexA). Ldh RNAi together with either human LDHA (UAS–hLdhA; D), human LDHB (UAS–hLdhB; E), or cytoplasmic LbNOX (UAS–LbNOX; F) expressed in surrounding glia (Repo-GAL4). Dashed outlines indicate progeny ($tdTomato^{+}$) derived from individual Dpn+ NSCs.

(G,H) EdU (NSC progeny, 6hr feeding), Dpn (NSCs) and tdTomato (NSC lineages) immunostaining in L3 VNCs in which either Control (LexAop-*mCherry*-RNAi; G) or Complex I (LexAop-*ND42*-RNAi; H) RNAi is expressed clonally in NSCs (Dpn-FRT-STOP-FRT-LexA). UAS-LbNox was overexpressed in surrounding glia (Repo-GAL4). Dashed outlines indicate progeny ($tdTomato^{+}$) derived from individual $Dpn^{+}$ NSCs.

(I) NSC proliferation (EdU+ progeny during a 6hr feeding period) in the VNC clonally expressing control or Complex I RNAi in NSCs while over-expressing UAS-LbNox in the surrounding glia. Each datapoint represents number of clones within individual brains, n = 78 clones, 16 brains [mosaic NSCs: Control RNAi; Glia: Control RNAi], 59 clones, 11 brains [mosaic NSCs: Control RNAi; Glia: LbNox], 34 clones, 7 brains [mosaic NSCs: Complex I RNAi; Glia: LbNox]; linear mixed effect models.

$p > 0.05$ = n.s.; $p < 0.05$ = *; $p < 0.01$ = **; $p < 0.001$ = ***. Scale bars are 5 µm (D-F,G,H).

**Supplementary Table 9. Drosophila RNAi lines for screening**

| REAGENT or RESOURCE | SOURCE | IDENTIFIER |
|---|---|---|
| UAS-Ldh-RNAi | BDSC | RRID:BDSC_33640 |
| UAS-Eaat1-RNAi | BDSC | RRID:BDSC_43287 |
| UAS-path-RNAi | BDSC | RRID:BDSC_64029 |
| UAS-hrm-RNAi | BDSC | RRID:BDSC_52902 |
| UAS-GOT1-RNAi | BDSC | RRID:BDSC_43194 |
| UAS-CG13384-RNAi | BDSC | RRID:BDSC_41703 |
| UAS-GOT2-RNAi | BDSC | RRID:BDSC_78778 |
| UAS-Gdh-RNAi | BDSC | RRID:BDSC_53255 |
| UAS-mAcon1-RNAi | BDSC | RRID:BDSC_34028 |
| UAS-CG43693-RNAi | BDSC | RRID:BDSC_35696 |
| UAS-Eaat2-RNAi | BDSC | RRID:BDSC_40832 |
| UAS-Idh-RNAi | BDSC | RRID:BDSC_41708 |
| UAS-Fatp-RNAi | BDSC | RRID:BDSC_55919 |
| UAS-CG1628-RNAi | BDSC | RRID:BDSC_54464 |
| UAS-Pcb-RNAi | BDSC | RRID:BDSC_56883 |
| UAS-aralar1-RNAi | BDSC | RRID:BDSC_56884 |
| UAS-CD98hc-RNAi | BDSC | RRID:BDSC_57746 |
| UAS-Gls-RNAi | BDSC | RRID:BDSC_62216 |
| UAS-GC1-RNAi | BDSC | RRID:BDSC_65937 |
| UAS-CG12773-RNAi | BDSC | RRID:BDSC_65949 |
| UAS-Balat-RNAi | BDSC | RRID:BDSC_67274 |
| UAS-Mpc1-RNAi | BDSC | RRID:BDSC_67817 |
| UAS-dmGlut-RNAi | BDSC | RRID:BDSC_36724 |
| UAS-Prestin-RNAi | BDSC | RRID:BDSC_50706 |
| UAS-Sfxn1-3-RNAi | BDSC | RRID:BDSC_38230 |
| UAS-slif-RNAi | BDSC | RRID:BDSC_64972 |
| UAS-CG8468-RNAi | VDRC | RRID:VDRC_6452 |

| | | |
|---|---|---|
| UAS-Indy-RNAi | VDRC | RRID:VDRC_9981 |
| UAS-CG10019-RNAi | VDRC | RRID: VDRC _7291 |
| UAS-WASp-RNAi | BDSC | RRID:BDSC_51802 |
| UAS-sns-RNAi | BDSC | RRID:BDSC_64872 |
| UAS-hbs-RNAi | BDSC | RRID:BDSC_57003 |
| UAS-Mct1-RNAi | VDRC | RRID: VDRC _106773 |
| UAS-SdhD-RNAi | BDSC | RRID:BDSC_65040 |
| UAS-Sln-RNAi | VDRC | RRID: VDRC_4607 |
| UAS-CG5535-RNAi | VDRC | RRID: VDRC_107030 |
| UAS-chaski-RNAi | VDRC | RRID: VDRC_37141 |
| UAS-kcc-RNAi | BDSC | RRID:BDSC_34584 |
| UAS-out-RNAi | BDSC | RRID:BDSC_67858 |
| UAS-CG4476-RNAi | BDSC | RRID:BDSC_38930 |
| UAS-Gs2-RNAi | BDSC | RRID:BDSC_40949 |
| UAS-PyK-RNAi | BDSC | RRID:BDSC_35218 |
| UAS-Mdh2-RNAi | BDSC | RRID:BDSC_36606 |
| UAS-Mdh1-RNAi | VDRC | RRID:VDRC_27398 |
| UAS-Alat-RNAi | BDSC | RRID:BDSC_60130 |
| UAS-Acly-RNAi | BDSC | RRID:BDSC_65175 |
| UAS-DGAT1-RNAi | VDRC | RRID:VDRC_100003 |
| UAS-Pld-RNAi | VDRC | RRID:VDRC_106137 |
| UAS-Lsd2-RNAi | VDRC | RRID:VDRC_102269 |
| UAS-Inx2-RNAi | BDSC | RRID:BDSC_29306 |
| UAS-sbm-RNAi | BDSC | RRID:BDSC_51791 |

**Supplementary Table 11. Primer sequences used for cloning**

| Primer name | Sequence (all 5'-3') |
|---|---|
| ATPsynO-shRNA-pFw | ctagcagtCTGGCTGACAACGGACGTCTAtagttatattcaagcataTAGACGTCCGTTGTCAGCCAGgcg |
| ATPsynO-shRNA-pR | aattcgcCTGGCTGACAACGGACGTCTAtagttatattcaagcataTAGACGTCCGTTGTCAGCCAGactg |
| ND-75-shRNA-pFw | ctagcagtCAAGGTGCTGTTCCTGTTGAAtagttatattcaagcataTTCAACAGGAACAGCACCTTGgcg |
| ND-75-shRNA-pR | aattcgcCAAGGTGCTGTTCCTGTTGAAtatgcttgaatataactaTTCAACAGGAACAGCACCTTGactg |
| ND-42-shRNA-pFw | ctagcagtTTGCTATGCTTTCGAGTTGAAtagttatattcaagcataTTCAACTCGAAAGCATAGCAAgcg |
| ND-42-shRNA-pR | aattcgcTTGCTATGCTTTCGAGTTGAAtatgcttgaatataactaTTCAACTCGAAAGCATAGCAAactg |
| mCherry-shRNA-pFw | ctagcagtCGAGTTCATCTACAAGGTGAAtagttatattcaagcataTTCACCTTGTAGATGAACTCGgcg |
| mCherry-shRNA-pR | aattcgcCGAGTTCATCTACAAGGTGAAtatgcttgaatataactaTTCACCTTGTAGATGAACTCGactg |
| UCP2-pFw | agggaattgggaattcgttaacagatctgcggccgcaaaatggttg |
| UCP2-pR | tctagattattacttgtcatcgtcatccttgtaatccatatggaag |
| Tra-NLS pF1 | ttgggaattcatgaagagcaggcacagaaggcatcgccagcgct |
| Tra-NLS pR1 | tttgcggccgcgtcttcgttcactgctgcgacttcggctccgttga |

**Supplementary Tables as separate files**

**Supplementary Table 1.** Provided as separate Excel file.
Single-cell RNA-seq cell-type marker genes.

**Supplementary Table 2.** Provided as separate Excel file.
EdgeR pseudo-bulk differential gene expression in each cell-type.

**Supplementary Table 3.** Provided as separate Excel file.
hdWGCNA module analysis on NSCs.

**Supplementary Table 4.** Provided as separate Excel file.
Gene Ontology analysis of NSC hdGWCNA modules.

**Supplementary Table 5.** Provided as separate Excel file.
Network analysis of NSC hdGWCNA modules.

**Supplementary Table 6.** Provided as separate Excel file.
Gene membership of NSC hdGWCNA modules.

**Supplementary Table 7.** Provided as separate Excel file.
Overlap of differentially expressed genes between edgeR and hdGWCNA in NSCs.

**Supplementary Table 8.** Provided as separate Excel file.
Transcription factor regulon analysis of differential gene expression in NSCs.

**Supplementary Table 10.** Provided as separate Excel file.
Larval brain metabolomics.
